# Mechanistic Studies of Small Molecule Ligands Selective to RNA Single G Bulges

**DOI:** 10.1101/2024.10.14.618236

**Authors:** Shalakha Hegde, Sana Akhter, Zhichao Tang, Chang Qi, Chenguang Yu, Anna Lewicka, Yu Liu, Kushal Koirala, Mikhail Reibarkh, Kevin P. Battaile, Anne Cooper, Scott Lovell, Erik D. Holmstrom, Xiao Wang, Joseph A. Piccirilli, Qi Gao, Yinglong Miao, Jingxin Wang

## Abstract

Small-molecule RNA binders have emerged as an important pharmacological modality. A profound understanding of the ligand selectivity, binding mode, and influential factors governing ligand engagement with RNA targets is the foundation for rational ligand design. Here, we report a novel class of coumarin derivatives exhibiting selective binding affinity towards single G RNA bulges. Harnessing the computational power of all-atom Gaussian accelerated Molecular Dynamics (GaMD) simulations, we unveiled a rare minor groove binding mode of the ligand with a key interaction between the coumarin moiety and the G bulge. This predicted binding mode is consistent with results obtained from structure-activity-relationship (SAR) studies and transverse relaxation measurements by NMR spectroscopy. We further generated 444 molecular descriptors from 69 coumarin derivatives and identified key contributors to the binding events, such as charge state and planarity, by lasso (least absolute shrinkage and selection operator) regression. Strikingly, small structure perturbations on these key contributors, such as the addition of a methyl group that disrupts the planarity of the ligand resulted in > 100-fold reduction in the binding affinity. Our work deepened the understanding of RNA-small molecule interactions and integrated a new generalizable platform for the rational design of selective small-molecule RNA binders.

## Introduction

RNA plays critical roles in gene regulation and various cellular processes in almost all life forms, including transcription, translation, splicing, and epigenetic modifications.^1,2^ Selective targeting of RNA structures using small molecules is an important pharmacological modality that complements traditional protein targeting approaches.^3–9^ For example, bacteria ribosomal RNA (rRNA) is an important antibiotic target with numerous clinically validated drug classes, such as aminoglycoside, tetracycline, macrolide, lincosamide, and oxazolidinone.^10^ Recently, two synthetic compounds, risdiplam and branaplam, both targeting precursor messenger RNA (pre-mRNA)-U1 small nuclear ribonucleoprotein (snRNP) complex, attracted tremendous attention as RNA splicing modulators to treat genetic diseases, including spinal muscular atrophy^11–17^ and Huntington’s disease.^18–20^ We previously demonstrated that a class of coumarin analogs of risdiplam can induce GA-rich sequences to form loop-like structures using molecular dynamics (MD) simulations and proposed that this interaction in cells provided additional selectivity of the coumarin derivatives to the GA-rich SMN2 gene.^21^

In addition to rRNA in bacteria and pre-mRNA in humans, several other classes of RNA have been targeted by chemical probes and drug candidates, including bacteria riboswitches,^22–24^ yeast self-splicing introns,^25^ microRNAs,^26–31^ untranslated regions of mRNAs^32–34^, and long non-coding RNAs (lncRNAs).^35–37^ In viruses, highly structured RNA regions have also been explored as targets for small molecules, such as an internal ribosome entry site (IRES) in the 5’-untranslated region (UTR) of the hepatitis C virus (HCV)^38–41^ and a transactivation response (TAR) hairpin in human immunodeficiency virus 1 (HIV-1).^42–44^ After the outbreak of SARS-CoV-2, we and others illustrated that the structural elements in the SARS-CoV-2 genome can also be targeted to suppress virus replication.^45–50^ Specifically, we discovered that some coumarin derivatives (**Fig. 1**) can be “repurposed” to selectively bind to a single G bulge in 5’ UTR of SARS-CoV-2 without retaining splicing modulatory activities or binding to GA-rich loops.^46^ We further demonstrated that covalently linking a ribonuclease (RNase) L recruiter and the coumarin-based G bulge binder yielded an active <u>ribo</u>nuclease <u>ta</u>rgeting <u>c</u>himera (RIBOTAC), which is effective in targeting SARS-CoV-2-infected epithelial cells.^45^

**Fig. 1.**
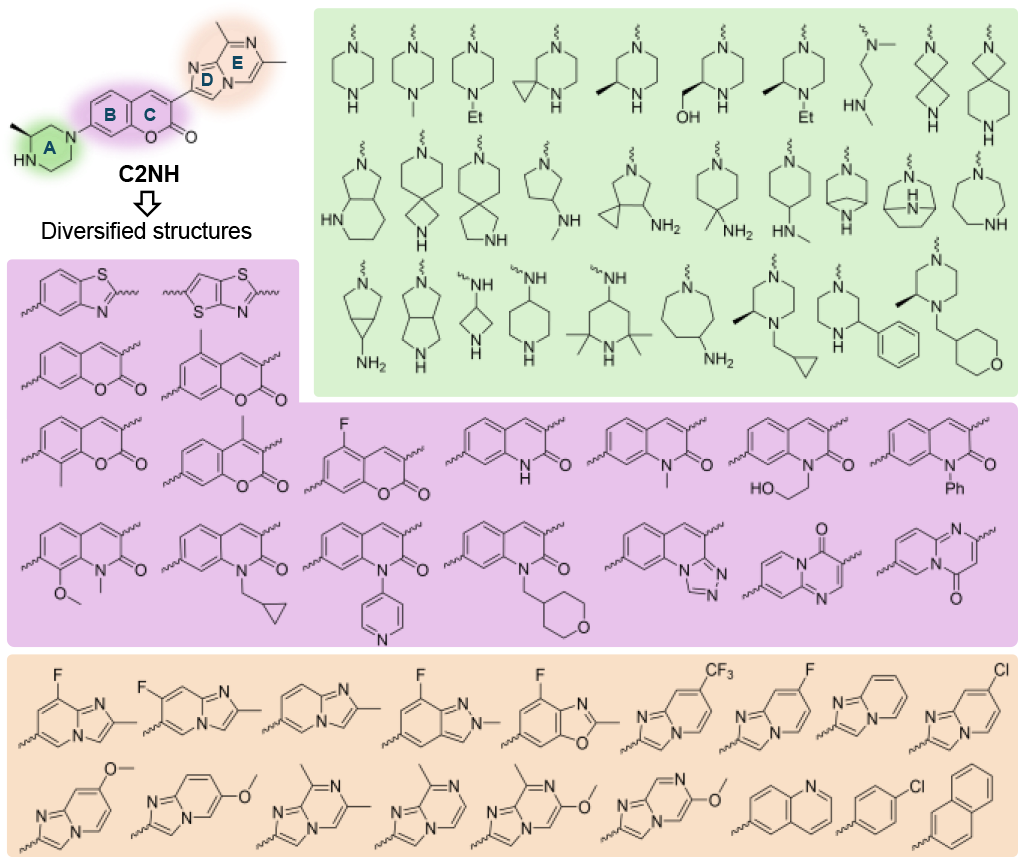
Molecular diversity of coumarin derivative analogs designed to bind bulged G RNA. The A, B/C, and D/E rings were replaced by the shaded green, purple, and orange structures, respectively (wavy lines = connecting bonds).

In general, attaining selective and effective targeting of RNA using small molecules is a challenging endeavor due to several factors, including its conformational flexibility/heterogeneity and its polyanionic backbone, which prevents the formation of deep hydrophobic binding pockets.^8,51^ Cheminformatics work has uncovered key factors governing the activity and selectivity of RNA binders^52–55^ and has been recently further advanced by machine learning approaches.^56^ However, the lack of methods for mechanistic studies of flexible RNA–small molecule ligand interactions critically limits further optimization of RNA ligands. A powerful approach to studying RNA-small molecule interactions is to use MD simulations,^21,57,58^ which are able to fully account for the RNA flexibility on an atomistic level. Here, we present an integrated platform that combines all-atom Gaussian accelerated MD (GaMD) simulations, which can rapidly predict ligand binding modes, with NMR transverse relaxation (R2) measurements and structure-activity relationship (SAR) studies that experimentally probe RNA–ligand interactions.^21,59^ We envision that our new platform for mechanistic studies on RNA ligands can provide a systematic approach to improving binding affinity for other RNA targets.

## Results and Discussion

### Coumarin derivatives selectively bind to RNA G (1×0) bulge

The prototype coumarin derivative C2NH, which binds to RNA single G bulges (denoted as G 1×0 bulge) at a moderate binding affinity, contains five heterocyclic rings: piperazine (A), coumarin (BC), and a [5,6]-fused ring (DE) (**Fig. 1**). We previously reported C2NH as an active splicing modulator that can bind to a GA-rich loop within the SMN2 gene.^21^ We modified the E ring to remove the splicing modulatory activity and repurposed the scaffold, resulting in a potent G 1×0 bulge binder that strongly associates with a structural motif in the RNA genome of SARS-CoV-2.^21,46^ To further probe the mechanism of the coumarin derivative in RNA binding interaction, we synthesized a collection of 69 analogs of C2NH (**Fig. 1**). Each compound in this collection comprises at least one ring distinct from the parent compound. For instance, in Ring A, the piperazine was replaced by cyclic amines of varying sizes. In Ring BC, the coumarin was substituted by other heterocycles with various substituents. Similarly, the [5,6]-fused Ring DE was replaced by [6,5]- or [6,6]-fused rings (**Fig. 1**).

All compounds in this collection are fluorescent with an excitation/ emission wavelength at ∼400/480 nm, which allowed us to use fluorescence polarization (FP) assay to rapidly determine their binding affinity to the bulge G RNA. Using a bulged G RNA segment from SARS-CoV-2 SL5 RNA (RNA1) as a model, we extensively profiled this 69-compound library against all four 1×0 RNA bulge variants (RNA1–4) for binding affinities (**Fig. 2A, Supplementary Fig. 1**, and **2**; **Supplementary Table1**). Almost all binding molecules showed superior selectivity for the G bulge compared to other RNA bulges (bulged A, U, and C), as judged by the polarization change (ΔmP) at two concentrations (1 and 5 μM) (**Fig. 2B**).

**Fig. 2.**
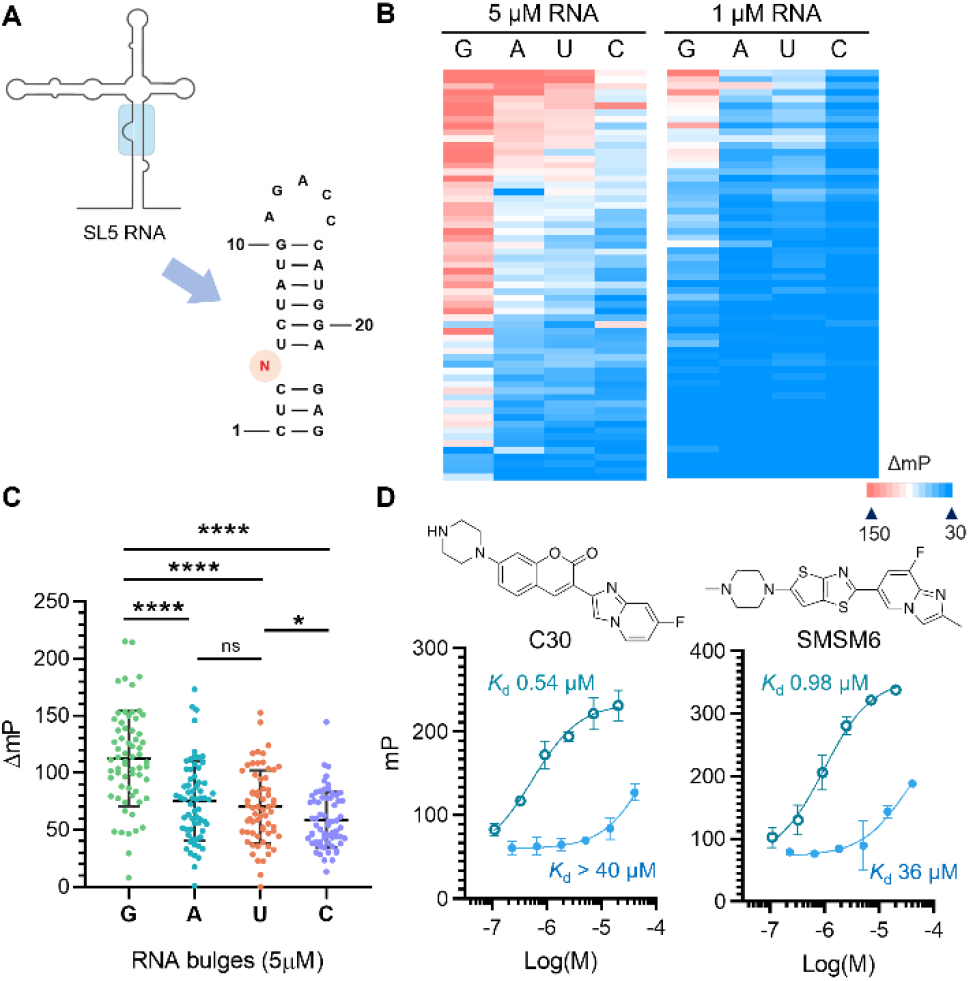
(A) RNA structures of the 1×0 RNA bulges used for in vitro binding profiling. N = G, A, U, or C (RNA1-4). (B) Heatmap profile of the ΔmP = (Polarization^RNA-ligand^ – Polarization^ligand^)×1000 for RNA binders in the presence of [RNA] = 5 or 1 μM (red = high polarization, blue = low polarization). (C) ΔmP of RNA-ligand complex for RNA ligands at 5 μM. Each data point represents a measurement of a ligand in the 69-compound collection. **** indicates p < 0.0001. (D) Dose-response curves for compounds selectively (SMSM6, C30) binding to bulged G RNA (RNA1) compared to an 11-nucleotide GA-rich sequence that would form a double loop-like RNA structure.

Statistical analysis of all 69 compounds revealed that the binding affinity for different RNA bulges followed the following trend: G >> A ≈ U > C (**Fig. 2C**). Since C2NH can also bind to GA-rich RNA loops^21^ via an induced-fit mechanism, resulting in the formation of a double loop-like structure, we tested the binding of coumarin derivatives to both a bulged G and a flexible GA-rich RNA (5’- U(GAAG)2GU). Interestingly, certain compounds, such as C30 and SMSM6, demonstrated > 35-fold selectivity towards bulged G over GA-rich RNA (**Fig. 2D**), while few compounds bind to both structures with comparable binding affinities (**Supplementary Fig. 3**). This suggests that coumarin derivatives may employ a distinct binding mechanism to selectively target RNA G bulges.

### GaMD simulations captured spontaneous minor groove binding of coumarin derivatives to RNA G (1×0) bulges

To explore the binding of specific RNA G bulge ligands, we performed all-atom simulations using the GaMD-enhanced sampling method.^59^ GaMD works by adding a harmonic boost potential to smooth the potential energy surface and reduce system energy barriers.^59^ GaMD has been shown to accelerate biomolecular simulations by orders of magnitude.^60,61^ For our GaMD simulations, we used the G bulge binder C30, which was among the most specific binders, and a model RNA hairpin with a G (1×0) bulge (RNA5: 5’-AAGAUGGAGAGCGAAACACACUCGUCUAUCUU; see **Extended Data Fig. 1A** for its secondary structure).

We found that C30 bound spontaneously to the G bulge and minor groove of RNA5 during the GaMD equilibration (**Extended Data Fig. 1B**) and three independent 1500 ns GaMD production simulations (**Fig. 3**). Upon binding to the minor groove of RNA5, the distance between the coumarin core of the ligand and the bulged G at position 24 (G24) was 3.5–5 Å, within a polar interaction range (**Fig. 3A**). Moreover, a π–π stacking interaction was observed between C12 and the fused D/E ring of the C30 ligand in simulations (**Fig. 3B**). We used these distances as reaction coordinates to further calculate a 2D free energy profile of C30 binding to RNA5, which showed two low-energy states, designated as “Bound” (more stable) and “Intermediate” states (**Fig. 3C**; Bound state structure was deposited in Model Archive Project ma-q6hl4). To experimentally probe the minor groove binding mechanism that we observed in the GaMD simulations, we conducted additional FP binding assays using C30 and various DNA versions of the RNA G bulge sequences (same sequences as RNA1 and RNA5). Our results show that the deoxyribose modification gives rise to a > 13-fold decrease in binding affinity (**Supplementary Fig. 4**). This result differed from what we observed with GA-rich loop binders, where the DNA aptamers bind to the ligands better than the RNA aptamers with the same sequences.^21^ Given that double-stranded (ds) DNA has a narrower minor groove than dsRNA,^62^ this result indicated that the groove region in dsDNA may not have sufficient space for C30 binding, supporting the minor grove binding mechanism.

**Fig. 3.**
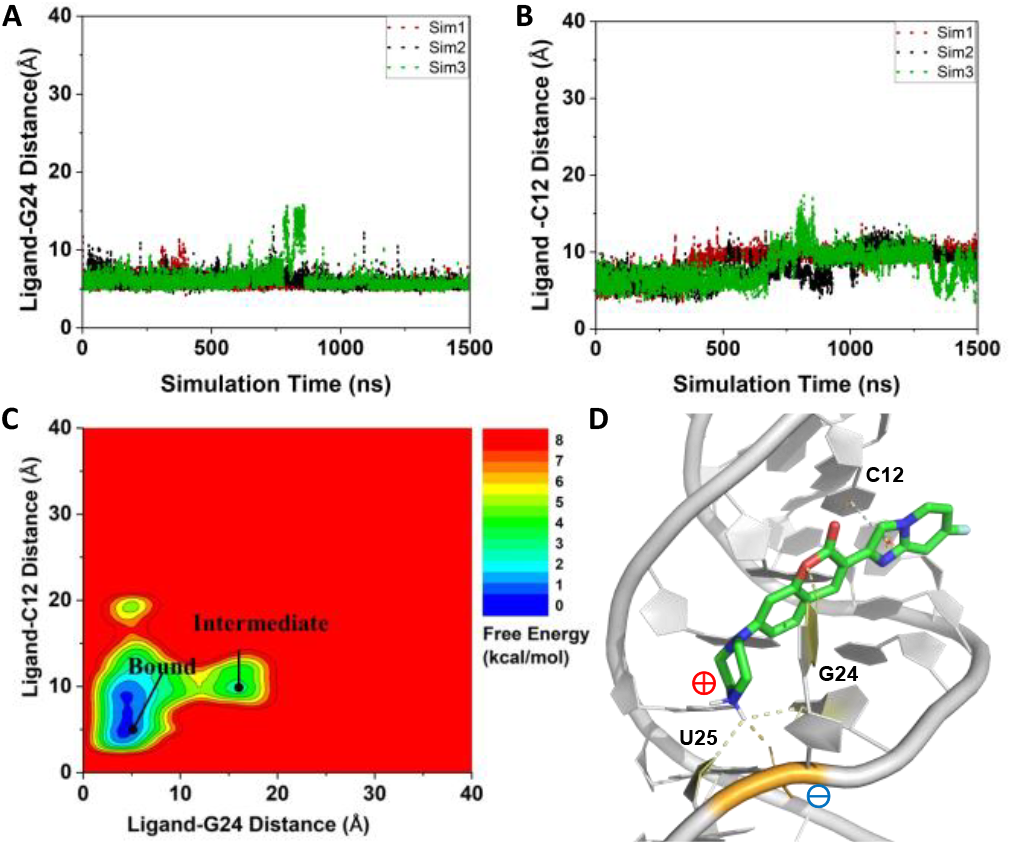
GaMD simulations captured stable binding of coumarin derivative C30 to RNA5. **(A)** Time courses of the center-of-mass distance between heavy atoms of the coumarin core in the ligand and the RNA bulge G24 calculated from three 1500 ns GaMD production simulations. **(B)** The center-of-mass distance between heavy atoms of the fused D/E ring of C30 and RNA nucleotide C12 plotted as a function of simulation time. **(C)** 2D free energy profile calculated with all three GaMD simulations combined, showing two distinct low-energy states, namely the “Bound” and “Intermediate”. **(D)** Representative conformation of RNA-C30 complex in the Bound state (grey dash line = π–π stacking, yellow dash line = hydrogen bonding, orange dash line = ionic interaction). The “Intermediate” conformation is shown in Extended Data **Fig. 2**.

In the simulation predicted “Bound” state, C30 formed three primary interactions within the minor groove of RNA5 (**Fig. 3D**). (1) The bulged G (G24) contributed to a hydrogen bond via its N1 position to the coumarin lactone moiety in C30. (2) A phosphate group in the RNA backbone was involved in an ionic interaction with the protonated NH2+ group in the piperazine ring of C30. (3) Nucleotide C12 formed π–π stacking interactions with the ligand C30 in the RNA minor groove. In the transient “Intermediate” state, C30 was located at a much larger distance from the G24 nucleotide and did not insert into the RNA minor groove (**Fig. 3C** and **Extended Data Fig. 2**).

We performed further GaMD simulations on two inactive analogs of C30, namely C30-Me (**Fig. 4A**) and SMSM64 (**Extended Data Fig. 3A**). C30-Me merely has an additional methyl group on the C ring compared to C30, which would break the planarity of Rings B/C and D/E in the compound (see discussions below). On the other hand, SMSM64 has an N-pyridinyl quinolone replacing the coumarin core, whose bulkiness might block the polar interaction with the RNA G bulge. In experiments, both compounds exhibited > 100-fold reduced binding affinities toward RNA5, with a dissociation constant (*K*d) of > 50 μM for both of the compounds, in comparison to C30, which has a *K*d of 0.27 ± 0.01 μM to RNA5 (**Supplementary Fig. 5**). Similar binding affinities were observed for these compounds when binding to RNA1 (see **Supplementary Table 1**).

**Fig. 4.**
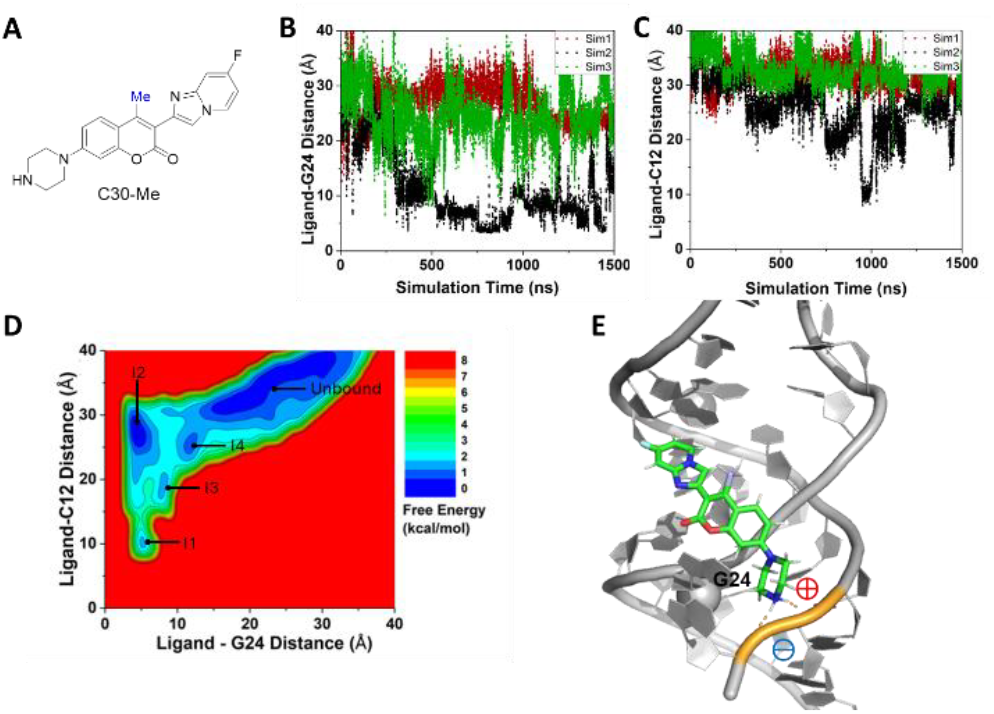
GaMD simulations captured transient binding of ligand C30-Me to RNA5: **(A)** Chemical structure of C30-Me. **(B)** Time courses of the center-of-mass distance between heavy atoms of the coumarin core in the ligand and the RNA bulge G24 calculated from three independent 1500 ns GaMD simulations. **(C)** The center-of-mass distance between the heavy atoms of the fused D/E ring of C30-Me and RNA nucleotide C12 plotted as a function of simulation time. **(D)** 2D free energy profile calculated with all three GaMD simulations combined, showing five low-energy states, namely the “I1”, “I2”, “I3”, “I4” and “Unbound”. **(E)** Representative conformation of C30-Me–RNA5 complex in the I1 state (orange dash line = ionic interaction).

In all three 1500 ns GaMD simulations for C30-Me, the ligand seldom reached the target site in the minor groove of RNA5 (**Fig. 4**). In the situation where C30-Me transiently interacted with G24 nucleotide (“Sim2” in **Fig. 4B**), the ligand remained out of the RNA minor groove with a distance > 10 Å from nucleotide C12 (**Fig. 4C**). Altogether, four transient binding states were identified from the free energy profile, designated as “Intermediate” states I1–I4, as well as the “Unbound” state, where the ligand dissociated from the RNA (**Fig. 4D** and **Extended Data Fig. 4**). The presence of multiple intermediate states suggested that the ligand explored various binding positions but was unable to achieve stable insertion into the minor groove. These intermediate conformations all maintained ionic interactions between the positively charged piperazine ring on the ligand and at least one phosphate group on the RNA backbone but could not fit into the minor groove (**Fig. 4E**). For SMSM64, the ligand remained mostly more than ∼15 Å away from key nucleotides G24 and C12 throughout the 1500 ns GaMD simulations (**Extended Data Fig. 3B** and **3C**). The resulting free energy profile showed only an “Unbound” state (**Extended Data Fig. 3D** and **3E**).

Interestingly, during the simulation studies, we observed a highly flexible G24 nucleotide within the RNA5 when the ligand was not bound to the RNA. Experimentally, we screened ∼20 crystal structures of RNA1 obtained using fragment antigen-binding region (Fab) chaperon-assisted crystallography (for a representative structure, see Protein Data Bank with accession code 9DN4) and observed dynamic conformations of the G bulge nucleotide,^63^ whereas other nucleotides remained relatively static (**Extended Data Fig. 5 and Supplementary Table 2**). This result is also consistent with our chemical probing results in SARS-CoV-2 RNA, where a high SHAPE (selecti e 2′ hydroxyl acylation analyzed by primer extension) signal was observed with high concentrations of acylating agents (e.g., 10 mM FAI-N3).^46^ Importantly, GaMD results demonstrated that the ligand with high affinity to RNA5 would bind and stabilize the flexible G bulge. Expectedly, we compared root-mean-square fluctuations (RMSFs) of each nucleotide in RNA5 across the simulated systems of C30, C30-Me, and SMSM64, and found that the interaction with C30 resulted in the lowest nucleotide G24 fluctuation throughout the simulation time course (**Extended Data Fig. 6**).

### NMR validation of the minor groove binding mode

Next, we used NMR experiments to validate the predicted binding mode between coumarin analog and bulged G RNA. First, we assigned imino protons and some other protons on the nucleobases in ^1^H NMR using a reported assignment that contains the segment of RNA5.^64^ The assigned peaks were distinguishable ones within 0.15 ppm from the reported ^1^H NMR chemical shifts (**Supplementary Table 3**). Next, we applied a recently published NMR method, ^1^H SOFAIR (band-Selective Optimized Flip-Angle Internally-encoded Relaxation),^65^ to quantify *R*_2_ relaxation rate of the receptor signals in order to characterize ligand-receptor interactions. *R*_2_ relaxation reflects on dynamics and motion changes of molecules, which is sensitive to weak binding (*K*d ∼*µ*M), and has been widely utilized as an NMR approach for identifying the binding sites of biomolecules^66,67^. Here, the SOFAIR pulse sequence^65^ was utilized to facilitate signal acquisition with high sensitivity of an RNA sample at mM concentration. Notably, SOFAIR was specifically designed to speed up data acquisition, and in this instance, led to a reduction in acquisition time from several hours, characteristic of conventional proton *R*_2_ measurements using Carr-Purcell-Meiboom-Gill (CPMG) type of methods,^68,69^ to ∼20 minutes.

The *R*_2_ relaxation rates were obtained from RNA nucleobases during the titration of the C30 ligand (**Supplementary Table 4**). As shown in **Fig. 5A**, the titration of C30 induced an overall *R*_2_ change, indicating binding between coumarin analog and bulged G RNA. The most pronounced increase in *R*_2_ was observed in G9, A10, U22, C23, and G24, implying the direct involvement of these nucleotides in binding. In contrast, the relaxation rates observed from G3 to G7 and C26 to C30 exhibited much smaller increases, or even negative changes upon ligand addition, suggesting that these regions of the RNA are not directly involved in binding. These findings from NMR experiments regarding the bound and unbound RNA nucleotides are consistent with those obtained from the GaMD simulations (**Fig. 5B**). Interestingly, the putative binding location is selective to one side of the G bulge (U22–G24), implying sequence selectivity in the minor groove. It is worth noting that only a few small molecular ligands have been reported as minor grove binders (e.g., PDB 1QD3),^70^ likely because the minor groove is wide and shallow in A-form dsRNAs. In our study, the NMR data strongly supported a minor groove binding mechanism for C30, as the ligand is unlikely to bind to the major groove of the RNA given the observed *R*_2_ relaxation changes. This finding further highlights the critical role of the bulged G in ligand interactions within this unusual minor groove binding mechanism.

**Fig. 5.**
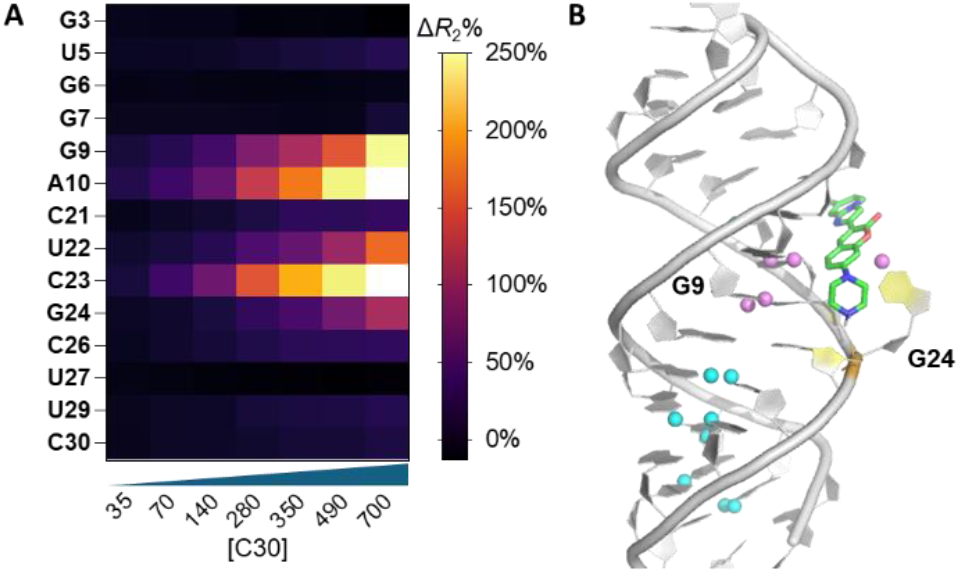
NMR relaxometry validation of the minor groove binding mode: **(A)** Normalized *R*_2_ relaxation rate percentage changes obtained from each assigned proton of RNA nucleotide in the absence and presence of C30 ligand. The colored columns in the bar plot represented *R*_2_ measured at different ligand concentrations. Normalized *R*_2_ values were calculated using the equation 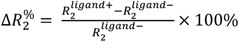 **(B)** Overlay of the simulation-predicted “Bound” state of C30-RNA5 and all identifiable protons in ^1^H SOFAIR (pink = increased Δ*R*_*2*_ ^%^; blue = unchanged or decreased Δ*R*_*2*_ ^%^).

### Molecular features on the ligands for RNA binding

We also determined how the molecular characteristics of these ligands contribute to their efficacy in RNA G bulge binding. Our approach involved a quantitative structure-activity-relationship (QSAR) investigation based on in vitro binding affinity data. Given the similarities in shape, size, and molecular scaffolds of our 69-compound library, we expected the QSAR analysis to offer detailed molecular insight into the specific structural and electronic properties responsible for their potency.

We used Molecular Operating Environment (MOE) software to individually predict the most likely protonation state based on the 2D structures of each molecule. The majority of molecules were found to be mono-protonated at the aliphatic cyclic amine (A ring), with a few exceptions that carried two positive charges (Supplementary Data). We then optimized the 3-dimensional (3D) structure of each molecule using ab initio density-functional theory (DFT) calculation with B3LYP 6-31G(d) basis set (Supplementary Data). Using these 3D structures as input, we generated 443 molecular descriptors using MOE software (Supplementary Data). To account for the planarity of the coumarin derivatives, we introduced a new dihedral descriptor between the plane B/C and D/E based on the most stable conformer predicted by DFT calculations. We then used the least absolute shrinkage and selection operator (lasso) regression technique to identify the important electronic and structural features among these molecular descriptors using a modified analytical pipeline.^56,71^ Lasso regression is a linear regression approach used for feature selection, which effectively eliminates unimportant variables. This process resulted in 16 molecular descriptors that significantly contributed to the binding affinity (**Supplementary Table 5**), of which eight molecular features are related to the charge and shape of the RNA ligands (**Table 1**).

**Table 1.**
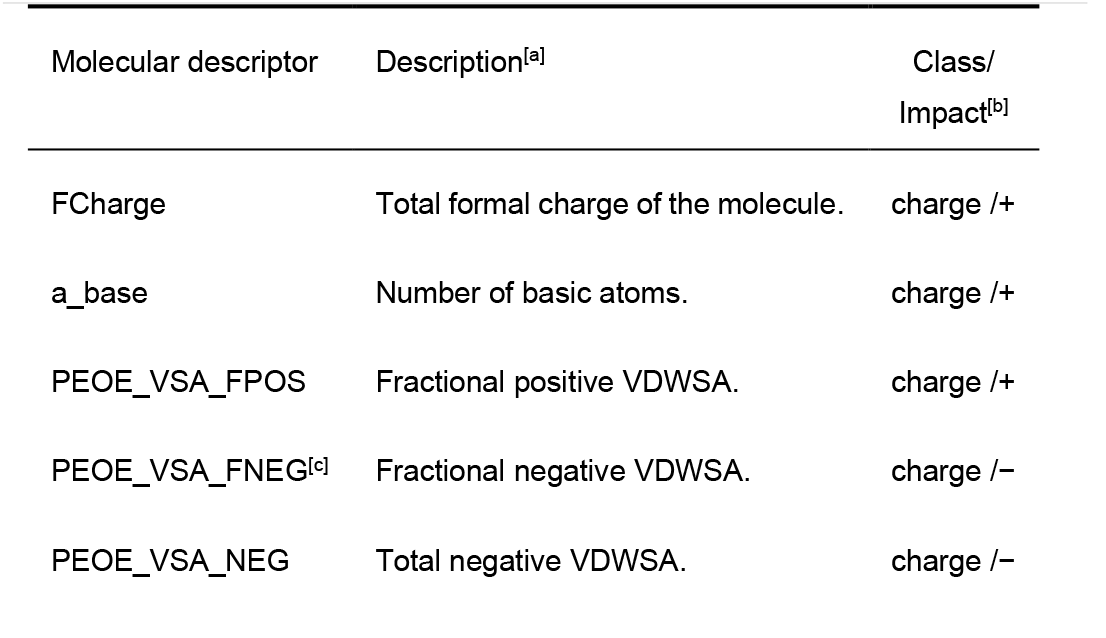
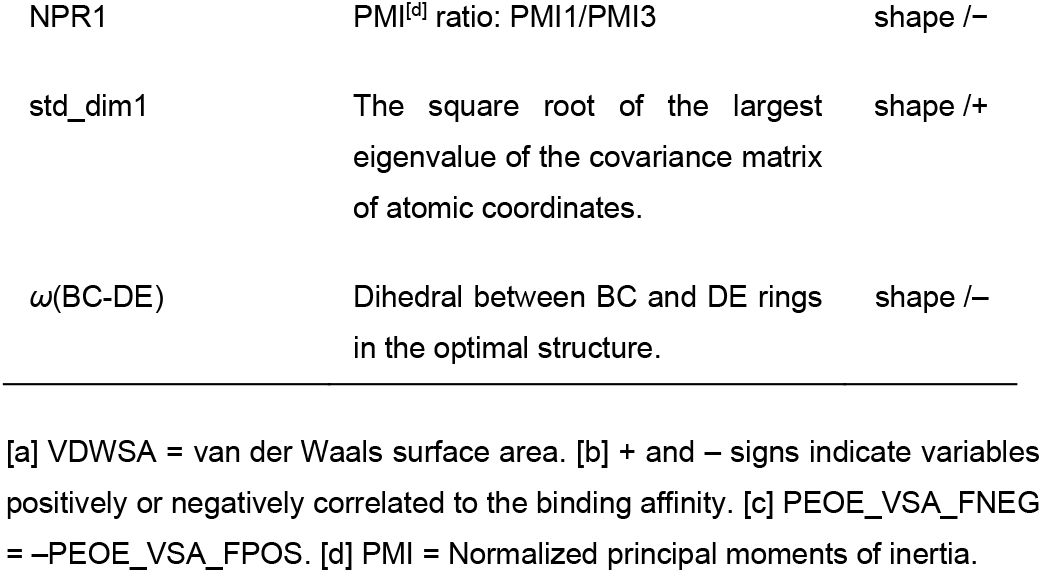
Molecular descriptors selected by lasso regression for RNA binding.^[a]^.

The five charge-related molecular descriptors were based on the total formal charge of the molecule (FCharge), the number of basic atoms that can be potentially protonated in physiological pH (A_base), the fractional positive (PEOE_VSA_FPOS) and negative (PEOE_VSA_FNEG) charges per unit area, and the total negative charge per unit area (PEOE_VSA_NEG). Since RNA is densely negatively charged, it is reasonable that positive charges would significantly contribute to RNA binding due to charge attraction. In the GaMD simulations with C30, intermolecular ionic interactions between the positive charge on the piperazine ring of C30 and phosphate groups on the RNA backbone were critical in maintaining the stability of the RNA-ligand complex. When we acetylated the piperazine ring of C30 at the N4 position (C30-Ac) to prevent protonation, the binding affinity decreased by a factor of > 5, highlighting the importance of electrostatic interaction between the ligand and RNA (**Supplementary Fig. 6**). We further hypothesized that the ligand used the positive charge on Ring A to explore suitable binding pockets at the early stage of the binding process. This hypothesis was supported by GaMD simulations, in which we observed all identifiable transient binding states (“Intermediate” states) of C30–RNA5 and C30-Me–RNA5 complexes retained an ionic interaction with RNA backbone phosphates (**Extended Data Fig. 2** and **Extended Data Fig. 4; Supplementary Movie 1**).

We also verified the impact of local positive charges on coumarin derivatives on in vitro binding by selecting four compounds, C29, C36, C34, and C34b, which only differ in the structures of the E ring. These compounds have two potential protonation sites: a piperazine A ring and an imidazole D ring. The second protonation site on the D ring can be partially stabilized by the coumarin moiety by forming an internal hydrogen bond. We speculated that the propensity of imidazolium formation significantly depends on the substituents on the E ring (**Fig. 6A**). For example, substituting the E ring with a trifluoromethyl group makes the molecule less amenable to protonation due to the electron-withdrawing effects. In contrast, the presence of an electron-donating methoxy group in compound C34b enhances the favorability of imidazolium formation. When the methoxy group is positioned at the 4’ location (C34), the existence of a resonance structure further contributes to stabilizing the positive charge (**Fig. 6A**). We verified the protonation energy of the four compounds relative to C29 using DFT calculations and compared it with the in vitro binding data (**Fig. 6B** and **6C**). The dissociation constants for these four compounds exhibit a consistent trend with respect to protonation energy, providing compelling evidence that local positive charges significantly contribute to the binding affinity.

**Fig. 6.**
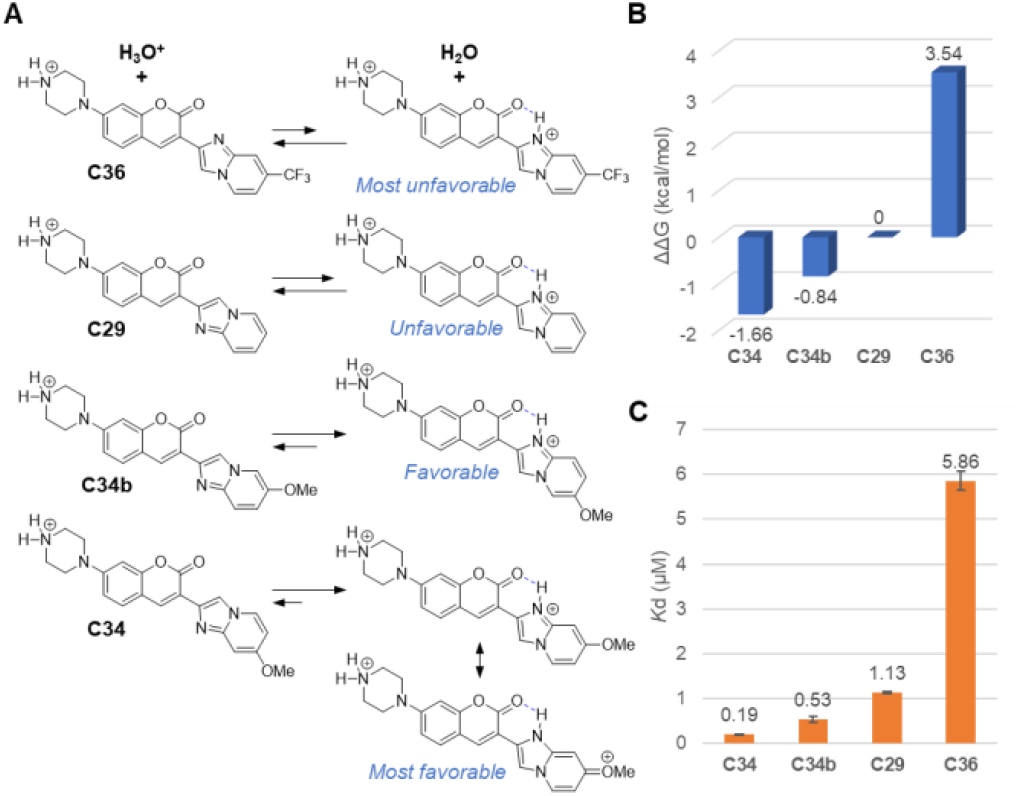
**(A)** Equilibria for the protonation reactions of four coumarin derivatives. **(B)** Protonation energy (relative to C29) was calculated using DFT with B3LYP 6-31G(d) basis set. **(C)** Observed binding affinity of the four compounds.

The 3D shape descriptors also strongly correlate with the binding affinity (**Table 1**). For example, NPR1 and NPR2 are numeric shape descriptors with values between 0 and 1 that characterize the general three-dimensional geometries of molecules.^72^ All compounds in our compound collection exhibit a small NPR1 value (< 0.2) and a large NPR2 value (> 0.85), indicating rod-like molecular structures. This observation is consistent with a prior cheminformatic analysis of diverse RNA-binding molecules.^53^ In addition, the positive contribution of the shape descriptor std_dim1 indicates that a longer molecule makes the ligand more favorable for binding, which is consistent with our expectations for groove binders.

Finally, we observed a positive correlation between planarity and binding affinity, as indicated by the inverse relationship between the dihedral angle of rings BC and E (ω(BC-DE)) and the natural logarithm of the binding constant (Ln*K*d). In C30, the dihedral angle between the BC-DE ring is ∼0, making it a planar molecule (**Extended Data Fig. 7**), which facilitates groove binding. However, adding a methyl group on ring C of C30 (C30-Me) causes steric hindrance between the methyl group and the lone pair electron of the imidazole nitrogen, disrupting the planarity of the molecule, rending it a poor binder (**Fig. 7A** and **Extended Data Fig. 7**). We also tested the role of this methyl group on ring B (C30-Me^RingB^), where it no longer sterically clashes with the imidazole ring. As expected, C30-Me^RingB^ is planar in its most favorable conformation (**Extended Data Fig. 7**), and the binding affinity was comparable to that of C30 (**Fig. 7A**). Planarity might also contribute to the high binding affinity of C34 (*Kd* = 0.10 ± 0.01 µM to RNA1). In the second protonation site of C34, the imidazole ring can form an internal hydrogen bonding with the coumarin lactone, further stabilizing the planar conformation (**Fig. 7B** and **Extended Data Fig. 7**).

**Fig. 7.**
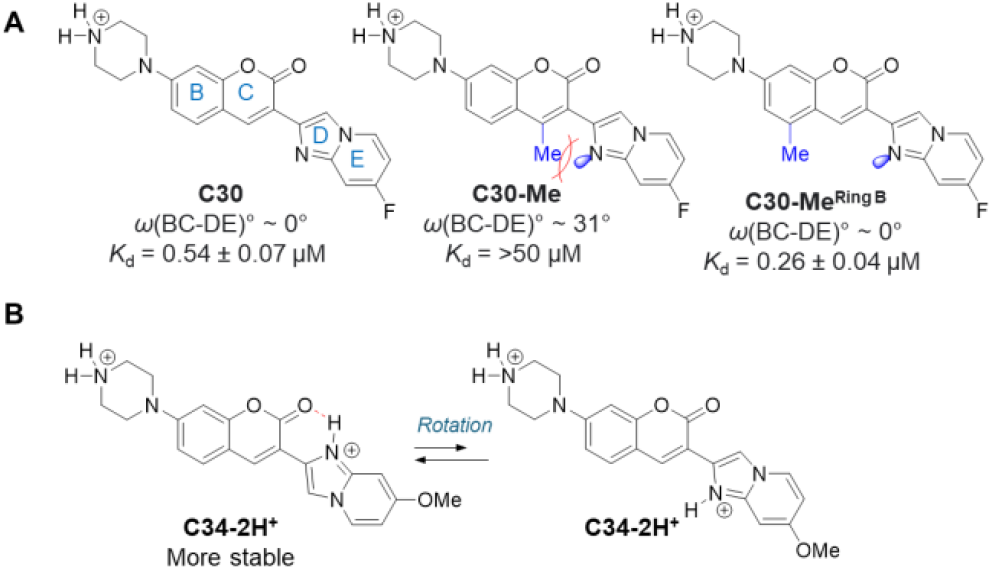
**(A)** Electronic clashes can be avoided by flipping the DE ring or forming an internal hydrogen bond between Rings C and D to maintain coplanarity. **(B)** C34 maintains planarity, favouring the in vitro interaction with Bulge G RNA.

## Conclusion

In this study, we have reported a new group of coumarin derivatives that exhibit selective binding to bulge G RNA. Using all-atom GaMD simulations, NMR, and QSAR studies, we have identified critical interactions that permit minor groove binding as well as crucial molecular properties of the ligands that significantly contribute to their binding affinity to bulge G RNAs. These factors include the shape and charge of the molecules. The minor groove ligand-RNA binding interface was validated by ^1^H SOFAIR NMR experiments that can rapidly characterize the RNA binding behavior. Our research represents a significant advancement in the understanding of RNA-small molecule interactions and offers a versatile platform for developing RNA binders with low nanomolar affinity.

## Methods

### Fluorescence polarization assay

Synthetic RNA oligomers were procured from GenScript (Piscataway, NJ, USA) and reconstituted in nuclease-free water. Compounds were prepared at a concentration of 50 µM in DMSO and were diluted in 2× assay buffer (50 mM MES, 100 mM NaCl, 0.004% TritonX, pH 6.1) to a concentration of 0.2 μM. A 1:3 dilution series (6 points) of each RNA was then prepared in 20– 30 μL water, resulting in concentrations ranging from 0.1–20 μM. Subsequently, 20 μL of 2× working solution containing the assay buffer and small-molecule ligand was added to each RNA sample in a 1:1 (v/v) ratio and mixed by pipetting. To measure fluorescence polarization, 8 μL of the resulting 1× sample solution was transferred into a 384-well, black, flat-bottom microplate (Greiner, #784076) in duplicates or triplicates. The plate was equilibrated at room temperature for 5 minutes prior to data acquisition using BioTek Synergy H1 (Winooski, VT, USA) with an excitation/emission of 360/460 nm at 25 °C. Experimental data were analyzed using the Prism 8.0 software package (GraphPad, San Diego, USA). A nonlinear curve fitting was employed to calculate the dissociation constant (*K*d), reported with a 95% confidence interval.

### Lasso regression

Lasso regression. The structure of each of the 69 compounds was individually optimized by DFT calculation with B3LYP/6-31G(d) basis set (ground state). The protonation state was predicted by MOE 2022 software (Chemical Computing Group, Montreal, Canada) at pH 7.0. The structures of all compounds were loaded on MOE 2022 and 443 molecular descriptors were generated using MOE 2022. Two molecular descriptors, h_pKa and h_pKb were excluded because the protonation states of the compound library are different. The dihedrals between BC and DE rings within the optimized 3-dimensional structure in the unit of degree (°) were added as a new molecular descriptor. In the lasso regression analysis, the natural logarithm of the dissociation constant (Ln(*K*d), *K*d in molar unit), expressed in molar units, was utilized as the dependent variable (y-value). (for final values of the molecular descriptors used for Lasso regression, see MolecularDescriptors.csv). Lasso regression (L1 regularization) was performed in R (4.2.2) according to the reported protocol.^71^ The non-zero coefficients were determined as described in **Supplementary Table 5**.

### Gaussian accelerated Molecular Dynamics (GaMD) Methodology

GaMD is an enhanced sampling technique in which a harmonic boost potential is added to smooth the potential energy surface and reduce the system energy barriers.^59^ GaMD can accelerate biomolecular simulations by order of magnitude and eliminates the need for predefined collective variables. Moreover, because GaMD boosts potential following a Gaussian distribution, it helps to properly recover biomolecular free energy profiles through cumulant expansion to the second order.^59^ A brief description of the method is described here.

Given a system with *N* atoms positioned at a specific location, 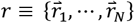, a boost potential 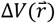 is added when the system potential energy 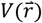 drops below the threshold energy *E*, the modified system potential 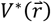 is calculated as:

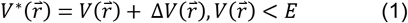

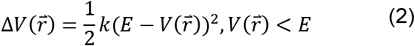

Where variable *k* represents the harmonic force constant. Two criteria must be met by the boost potential 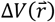. First, for any two arbitrary potential values 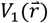 and 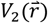 found on the original energy surface, if 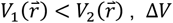 should be a monotonic function that does not change the relative order of the biased potential values 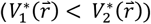. Second, if 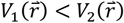, the potential difference observed on the smoothened energy surface should be smaller than that of the original 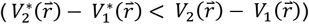. By combining the first two criteria and plugging in *1* and *2*, we obtain:

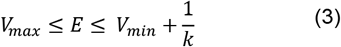

where *V*_*min*_ and *V*_*max*_ are the system minimum and maximum potential energies and *k* satisfies: 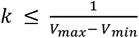. If we define 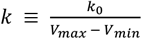, then 0 < *k*_0_ ≤ 1. The greater the *k*_0_ value is, the higher the boost potential 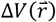 is added to the potential energy surface. Third, the standard deviation of Δ*V* should be small (i.e., narrow distribution) for accurate reweighting using cumulant expansion to the second order:

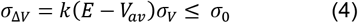

in which *V*_*av*_ and *σ*_*V*_ are the average and standard deviation of the system potential energies and *σ*_Δ*V*_ is the standard deviation of Δ*V* with *σ*_0_ as a user-specified upper limit (10*k*_*B*_*T*) for precise reweighting. Eq. 3 states that, when *E* is set to the lower bound *E* = *V*_*max*_, *k*_0_ can be calculated as:

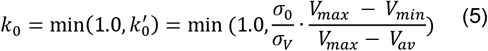

On the other hand, when the threshold energy *E* is set to its upper bound 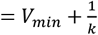, *k*_0_ is set to:

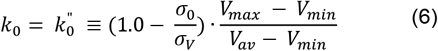

if 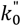 is calculated between 0 and 1. Otherwise, *k*_0_ is calculated using Eq 5.

The GaMD method provides options to add only the total potential boost Δ*V*_*p*_, only the dihedral potential boost Δ*V*_*D*_, or the dual boost potential (both Δ*V*_*p*_ and Δ*V*_*D*_). Among these, the dual-boost GaMD (GaMD_Dual) mode provides the highest acceleration for enhanced sampling of simulations.

The simulation parameters comprise the threshold energy *E* for applying boost potential and the effective harmonic force constants, *k*_0*p*_ and *k*_0*D*_ for the total and dihedral boost potential, respectively.

Example input parameters used in dual-boost GaMD simulations include the following in addition to those used in conventional MD: igamd = 3, iE = 1, irest_gamd = 0, ntcmd= 1500000, nteb = 30000000, ntave = 300000, ntcmdprep = 600000, ntebprep = 600000, sigma0P = 6.0, sigma0D = 6.0

### Energetic Reweighting of GaMD simulations

In biomolecular systems, the probability distribution along a selected reaction coordinate A(r) is represented as *p*(A)*, where r is representative of the atomic locations. The canonical ensemble distribution, *p(A)*, can be recovered by reweighting *p*(A)* using each frame’s boost potential, *ΔV(r)*.

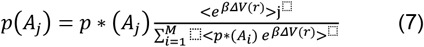

where *M* is the number of bins, *β* = *k*_*B*_*T* and 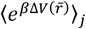 is the ensemble-averaged Boltzmann factor of 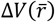 for the simulation frames found in the *j*^th^ bin. To approximate the ensemble-averaged reweighting factor, one can use the cumulant expansion method.:

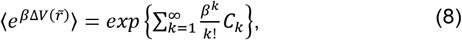

where the first two cumulants are given by:

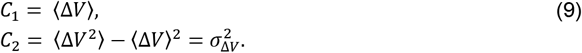

The boost potential obtained from GaMD simulations shows a near-Gaussian distribution. Cumulant expansion to the second order thus provides a good approximation for computing the reweighting factor. The reweighted free energy *F*(*A*) = −*k*_*B*_*T* In *p*(*A*) is calculated as:

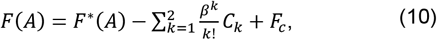

where *F*^∗^(*A*) = −*k*_*B*_*T* In *p*^∗^(*A*) is the modified free energy obtained from GaMD simulation and *F*_*c*_ is a constant.

### NMR experiments

An 84.1 nmol unlabeled RNA5 sample (Sigma-Aldrich) was dissolved in 135 *µ*L of potassium buffer (25 mM potassium phosphate buffer, 50 mM potassium chloride, pH 6.2, 10% D2O) to prepare an ∼600 *µ*M NMR sample. A similar sample condition was used in a previous report,^64^ where NMR assignment was determined for a ^15^N-labeled RNA containing the segment of RNA5.

All NMR spectra were acquired using a Bruker 800 MHz Ascend spectrometer equipped with a TCI cryoprobe at 298 K **(Extended Data Fig. 8**). RNA proton peak assignment was performed by comparing the measured ^1^H chemical shifts with literature values (**Supplementary Table 3**).^64^ Proton peaks were assigned if the chemical shift difference was less than 0.15 ppm. All NH protons of U and G residues that showed peaks in the 10-14 ppm region were assigned accordingly, except for the one from G24, where the literature assignment was missing. Two proton peaks observed in this chemical shift region that were not previously assigned should correspond to the NH of G24 as well as that of G13. G13 is a part of the linker that differs from the RNA sequence used in the literature. The proton peaks of G7-NH and U25-NH did not appear until the addition of a 70 *µ*M ligand. All the resonances that were unambiguously assigned are summarized in **Supplementary Table 3**.

The ligand stock solution prepared for the titration experiment contained 10 mM C30 in DMSO-d6. 0.47 *µ*L, 0.47 *µ*L, 0.94 *µ*L, 1.88 *µ*L, 0.94 *µ*L, 1.88 *µ*L, and 2.82 *µ*L of the stock solution were titrated into the RNA NMR sample to achieve final ligand concentrations of 35 *µ*M, 70 *µ*M, 140 *µ*M, 280 *µ*M, 350 *µ*M, 490 *µ*M, and 700 *µ*M, respectively. Proton transverse relaxation rate *R*_2_ was measured from SOFAIR (band-selective optimized flip-angle internally-encoded relaxation) (**Extended Data Fig. 9**).^73^ The band-selective excitation pulse p39 was centered at 12.1 ppm with a bandwidth of 5.2 ppm for NH region, and was centered at 7.6 ppm with a bandwidth of 3 ppm for NH2/aromatic region. Transverse relaxation was encoded through the incrementation of a delay *t* flanking the refocusing pulse p40. The delay time was set to between 0 to 0.4 s with a total of 12 increments (0, 0.002, 0.005, 0.010, 0.015, 0.020, 0.025, 0.030, 0.050, 0.1, 0.2, 0.4 s). The duration of each experiment is about 18.5 minutes. Data were processed and analyzed using MestReNova. *R*_2_ of each resonance was determined through area integration and fitting the integrals to the following equation: 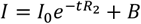, where *t* is the delay time, *R*_2_ is the transverse relaxation rate, and *B* is a constant to account for any baseline differences between experiments.

## Data Availability Statement

The representative C30-RNA5 bound conformation generated by GaMD simulations is available in PDB format in the Model Archive repository (https://modelarchive.org) under project ma-q6hl4. The Representative apo RNA crystal structure of RNA1 is available in PDB under accession number 9DN4.

## Supporting Information

Supplementary figures and tables, experimental and computation methods, and compound characterization data are available in the Supporting Information file. Calculated molecular descriptors (csv format), the code used for lasso regression (R markdown file), optimized structures for the 69-compound collection (compressed mol2 file), and simulated C30-RNA5 binding pathway (Supplementary Movie 1, mp4 movie) are also available in the Supporting Information.

## Acknowledgements

Research reported in this article was supported by the National Institute of General Medical Sciences (NIGMS) of the National Institutes of Health (NIH) under award numbers R35GM147498 (to J.W.) and start-up project 27110 at the University of North Carolina – Chapel Hill (to Y.M.). The authors acknowledge support from the MRL Postdoctoral Research Program, and Xingjian Xu at Merck & Co., Inc., Rahway, NJ, USA, for discussions of NMR measurements. This research used the NYX beamline 19-ID, supported by the New York Structural Biology Center, at the National Synchrotron Light Source II, a U.S. Department of Energy (DOE) Office of Science User Facility operated for the DOE Office of Science by Brookhaven National Laboratory under Contract No. DE-SC0012704. The NYX detector instrumentation was supported by grant S10OD030394 through the Office of the Director of the National Institutes of Health. The Center for Bio-Molecular Structure (CBMS) is primarily supported by the NIH-NIGMS through a Center Core P30 Grant (P30GM133893), and by the DOE Office of Biological and Environmental Research (KP1607011). NSLS2 is a U.S.DOE Office of Science User Facility operated under Contract No. DE-SC0012704.This publication resulted from the data collected using the beamtime obtained through NECAT BAG proposal # 311950.

## Extended data

**Extended Data Fig 1:**
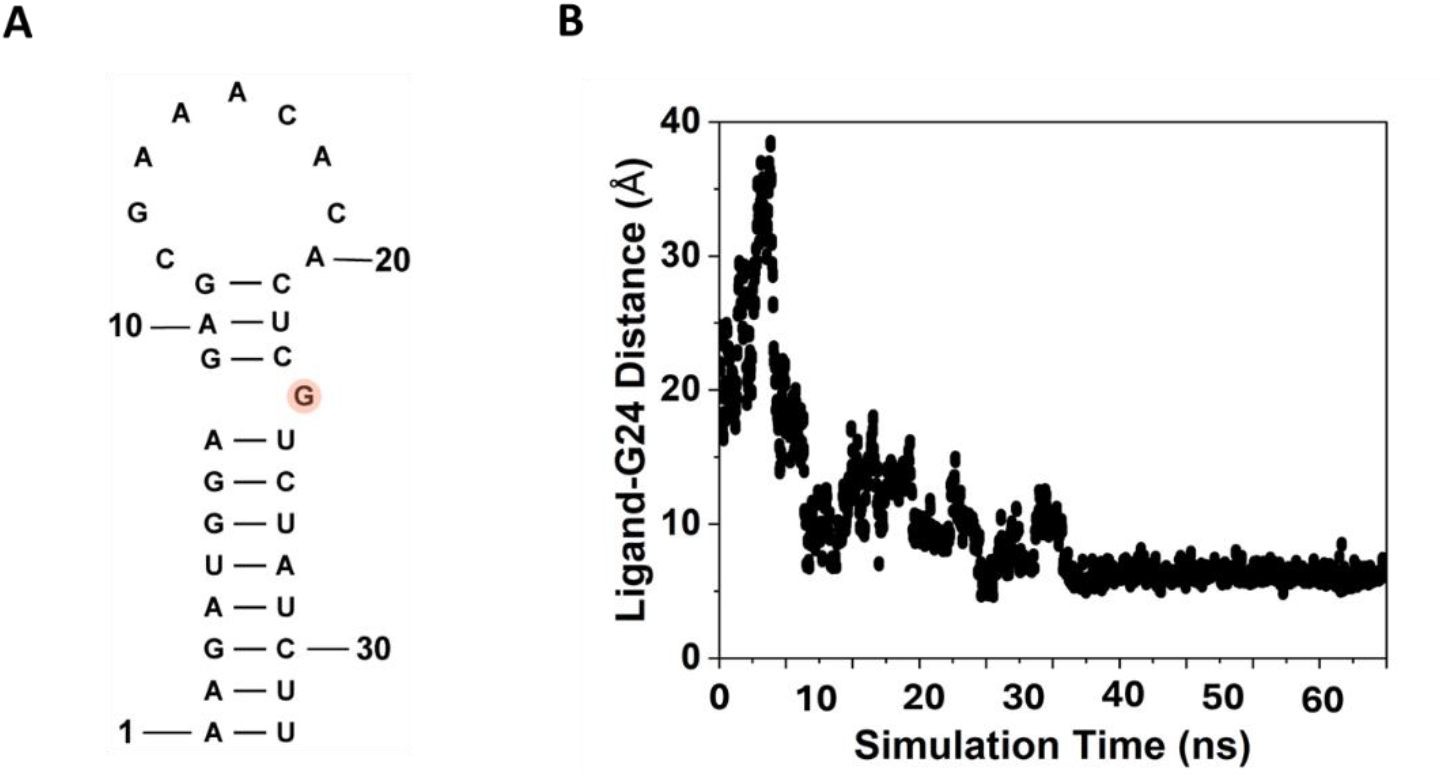
Binding of C30 with RNA5: **(A)** Structure of RNA5. **(B)** Time courses of the center-of-mass distance between the ligand-heavy atoms in the coumarin core to the RNA bulge G (RNA5) calculated from 63 ns GaMD equilibration simulation.

**Extended Data Fig. 2:**
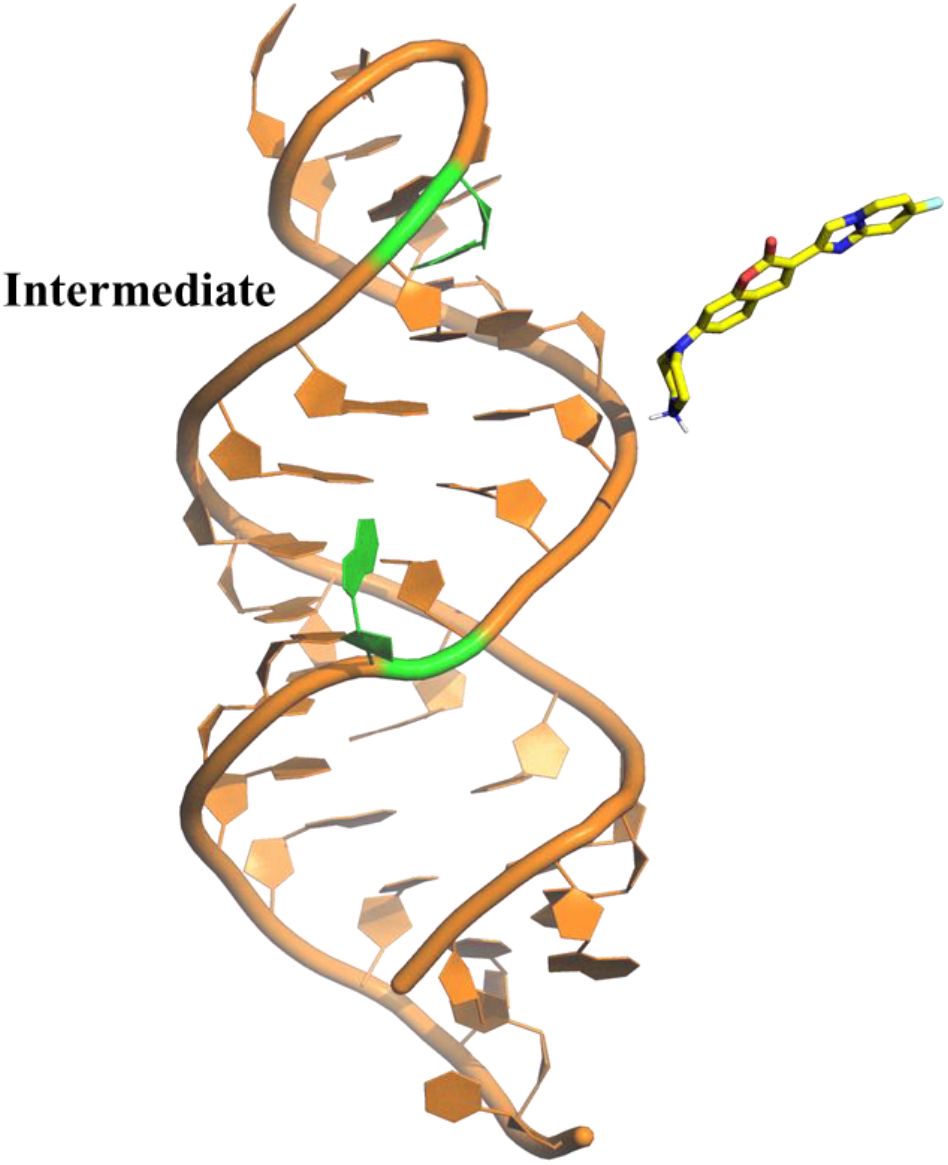
The “Intermediate” state of RNA5−C30 binding obtained from the GaMD simulations.

**Extended Data Fig. 3:**
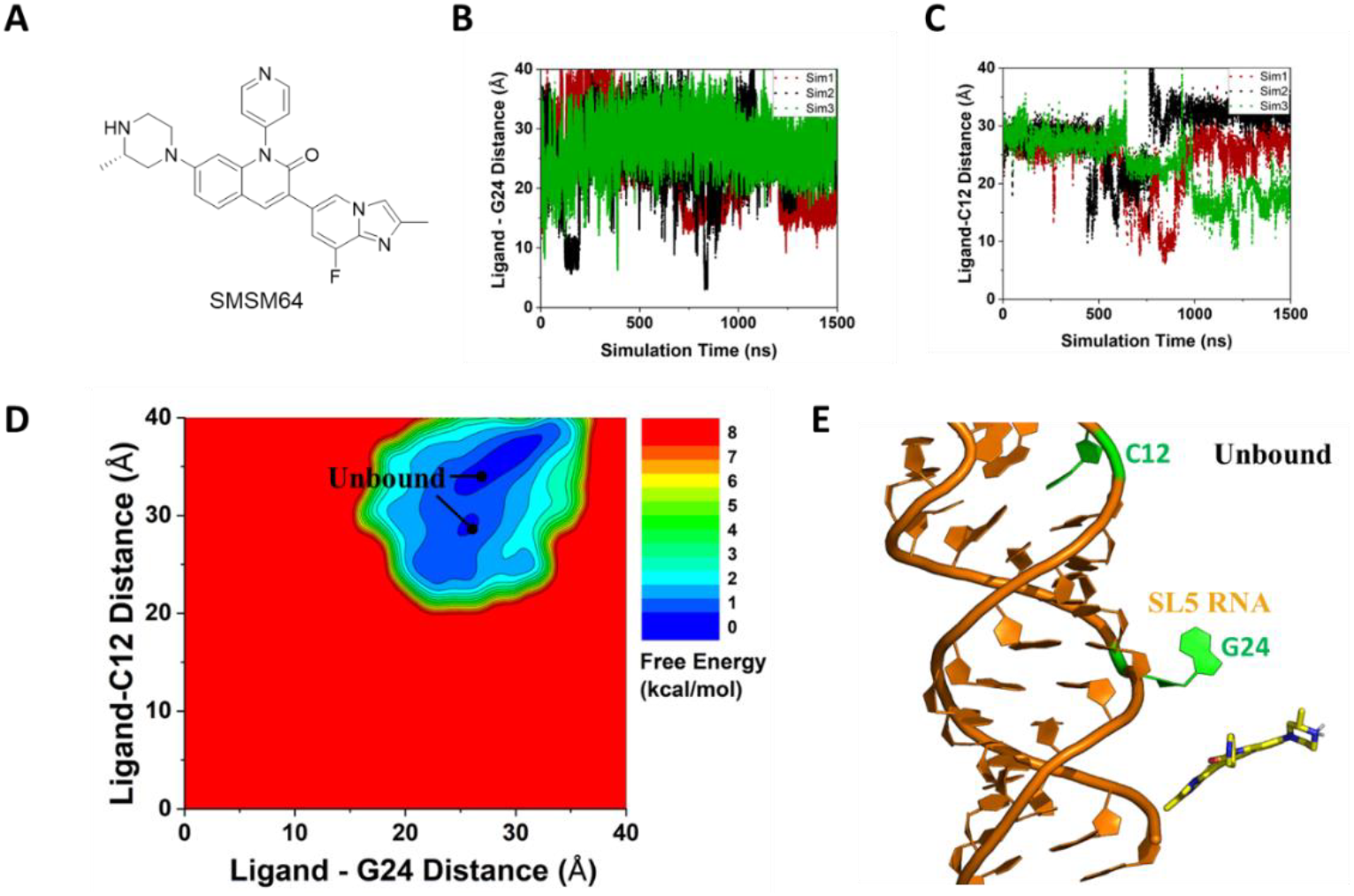
GaMD simulations captured spontaneous binding of coumarin derivative SMSM64 to RNA5: **(A)** Time courses of the center-of-mass distance between ligand heavy atoms in the coumarin core to the RNA bulge G24 calculated from three independent 1.5 µs GaMD simulations. **(B)** π–π stacking interaction distance between the fused (D, E) ring and nucleotide C12 tracked as a function of simulation time. **(C)** 2D free energy profile calculated with all three GaMD simulations combined, in which a single distinct low-energy state was identified, referred as the “Unbound” state. **(D)** Representative conformation of RNA not bound to SMSM64.

**Extended Data Fig. 4:**
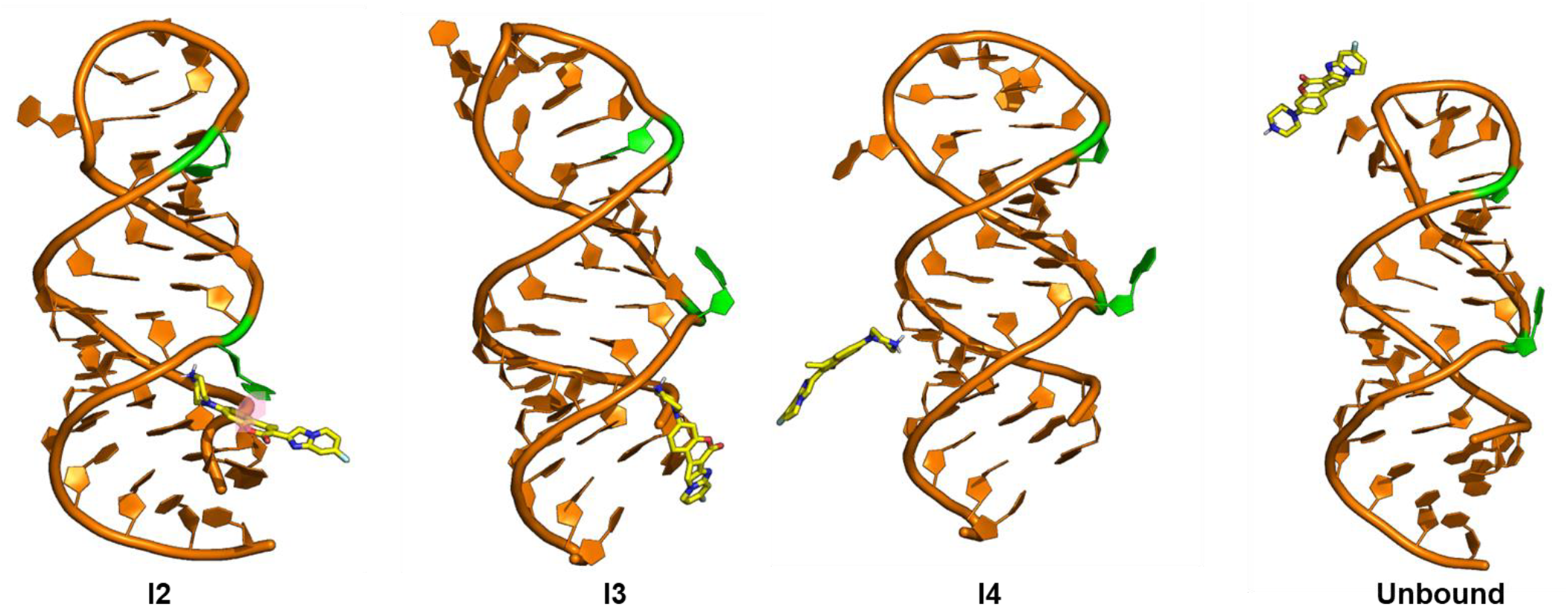
Intermediate conformational states of C30-Me binding to the RNA5. Intermediate states as identified from the 2D free energy profile from GaMD simulations, “I1”, “I2”, “I3”, “I4” and “Unbound” states. The RNA is in orange cartoons, and the ligand is in sticks with the C atoms, which are colored yellow. The unpaired G24 and C12 nucleotides are highlighted in light green.

**Extended Data Fig. 5:**
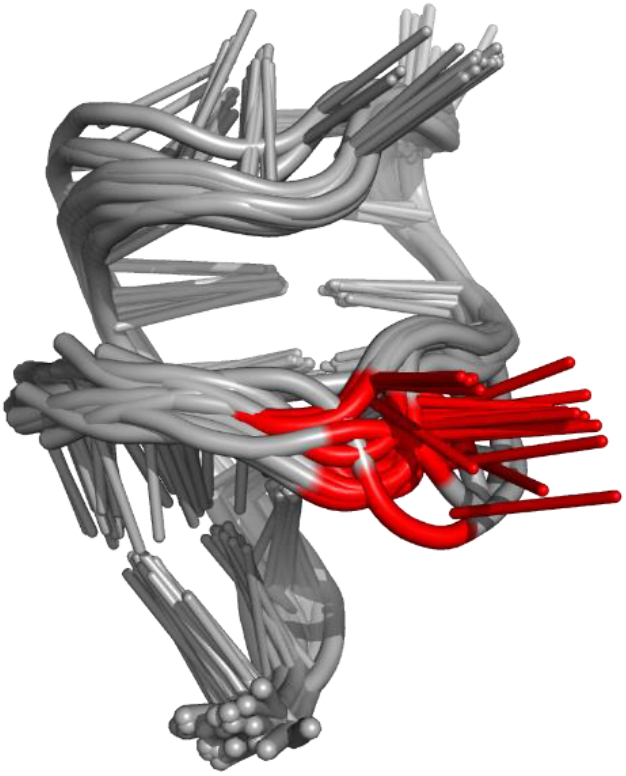
Chaperone-assisted crystal structure of RNA 1. Structural superposition of 21 RNA molecules derived from 12 datasets. Among aligned structures, 9 datasets consist of two molecules, and 3 datasets consist of one molecule. All models were refined at least 3 times. Resolution of collected datasets ranges from 1.39 Å to 2.45 Å; completeness from 93 % to 100 %. Structures were solved in triclinic, monoclinic and orthorhombic crystal systems.

**Extended Data Fig. 6:**
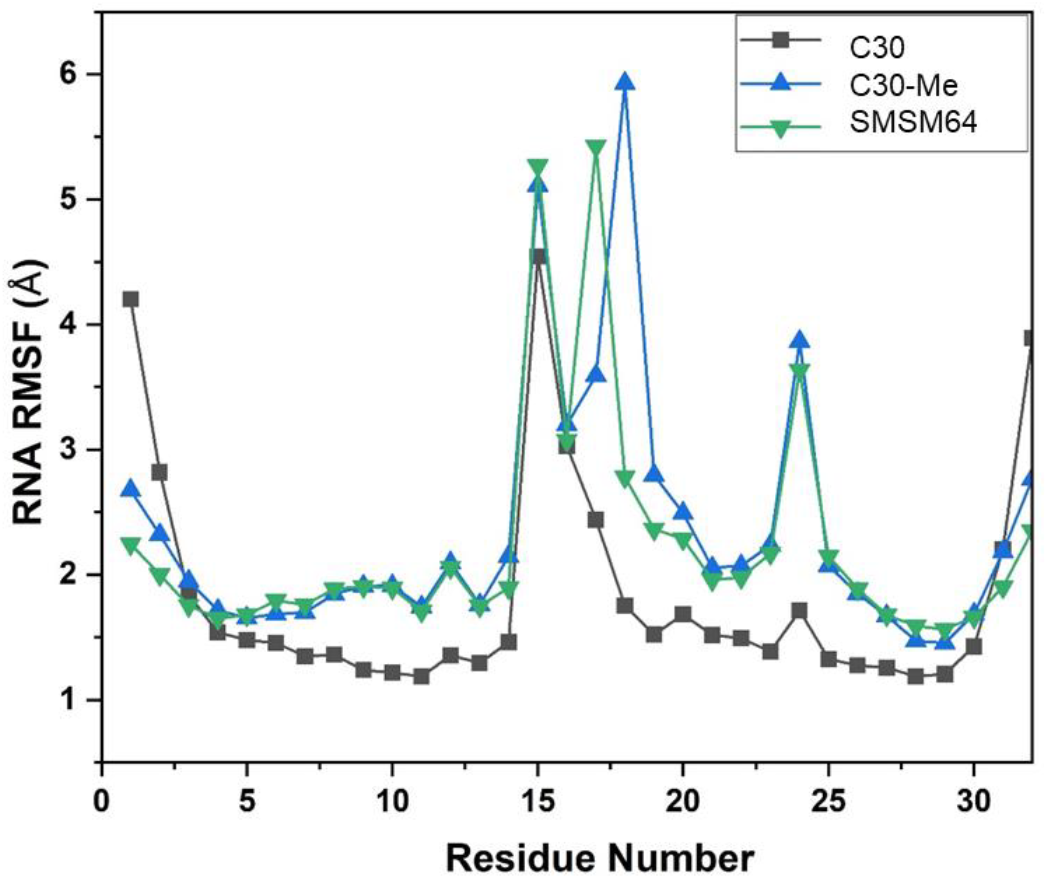
Fluctuation of the RNA nucleotides across GaMD production simulation in the presence of the coumarin derivatives. RMSF = Root Mean Square Fluctuation.

**Extended Data Fig. S7:**
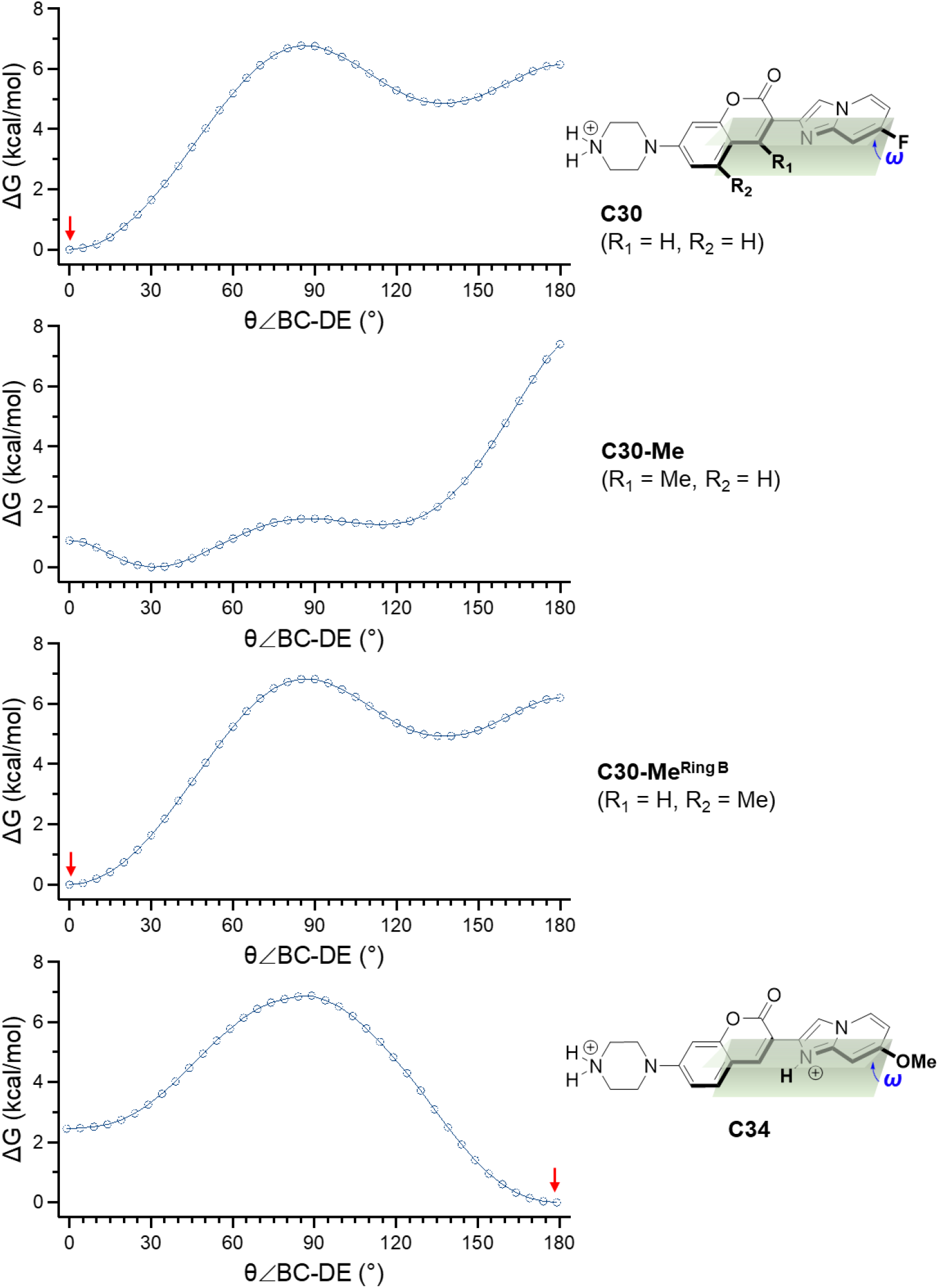
Calculated total energy of coumarin derivatives with a scan of ω(BC-DE) dihedral (0–180°). The calculation was performed using DFT with a B3YLP 6-31(d) basis set. The red arrow indicates the most stable planar structures.

**Extended Data Fig. S8:**
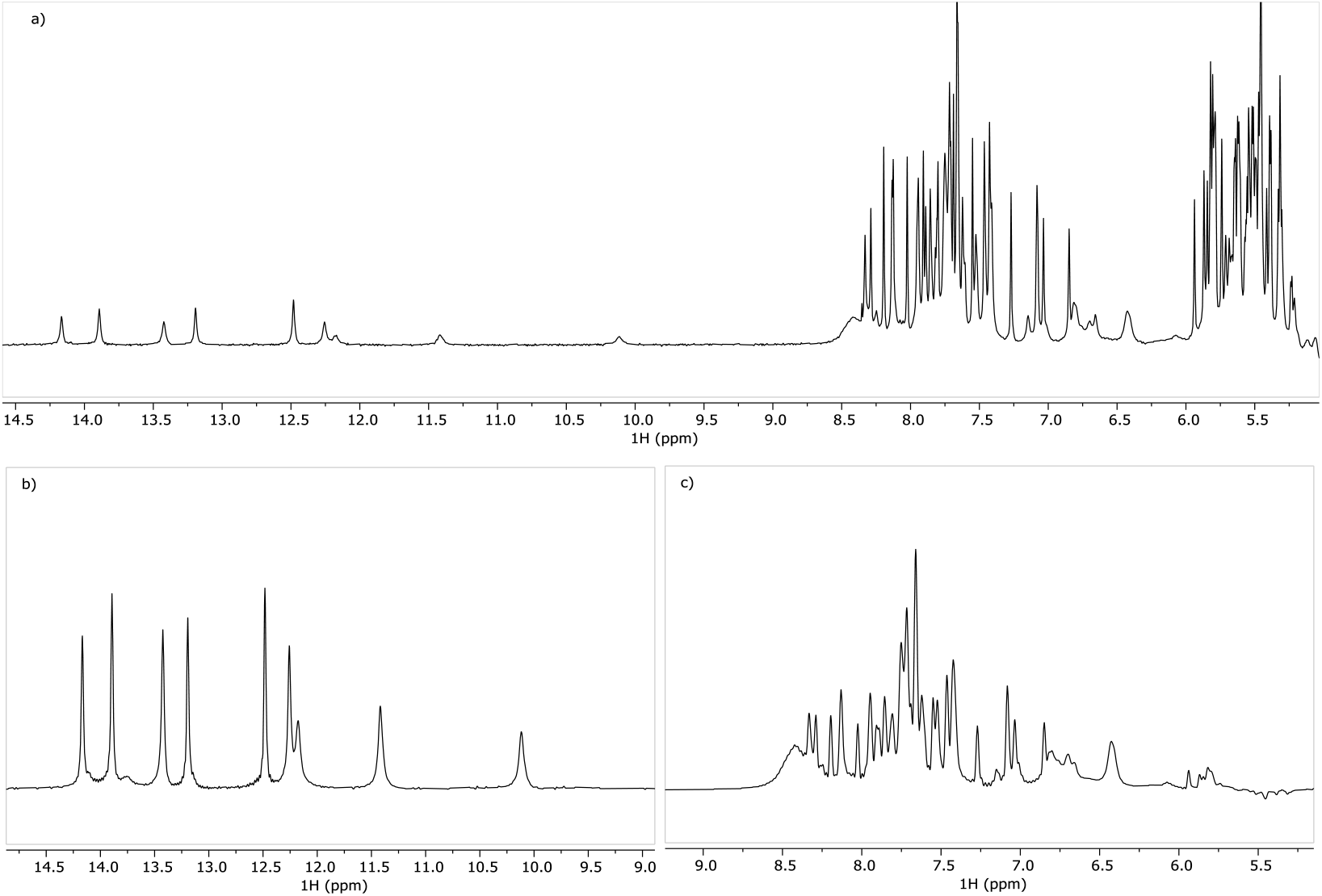
NMR spectra of RNA5. **(A)** 1D ^1^H experiment with solvent suppression. **(B)** The first increment of SOFAIR experiment with a center at 12.1 ppm. **(C)** The first increment of SOFAIR experiment with chemical shift center at 7.6 ppm.

**Extended Data Fig. S9:**
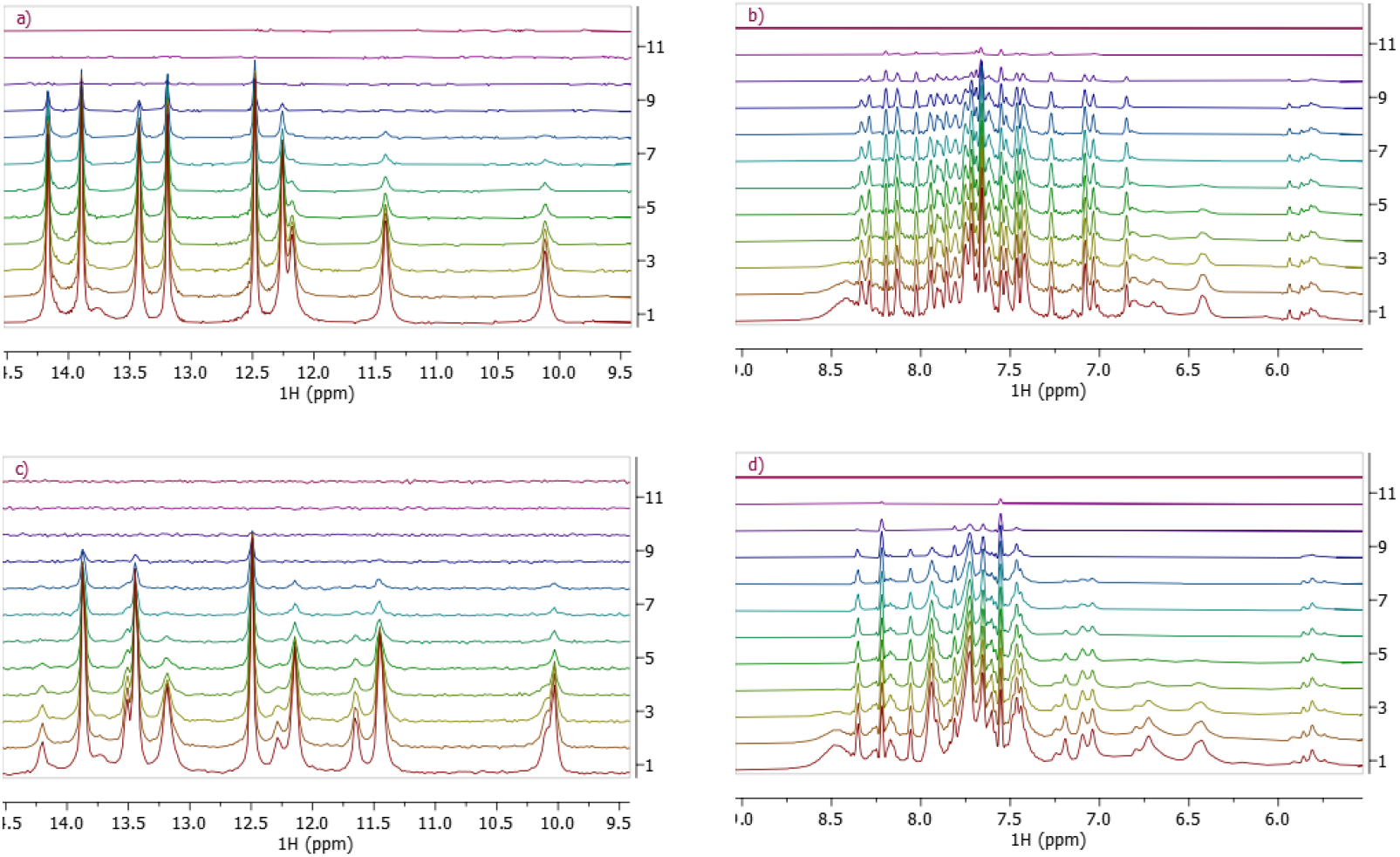
R2 relaxation measurement of RNA5 using SOFAIR experiments. **(A)** and **(C)** measured R2 relaxation rates in the range of 10-14 ppm without and with 700 µM C30 ligand. **(B)** and **(D)** measured R2 relaxation rates in the range of 6-8 ppm without and with 700 µM C30 ligand. Vertical spectra were obtained with spin echo intervals of 0, 0.002, 0.005, 0.01, 0.015, 0.02, 0.025, 0.03, 0.05, 0.1, 0.2, and 0.4 s from bottom to top.

