## Supporting Information for "Mechanistic Studies of Small Molecule Ligands Selective to RNA Single G Bulges"

### Abbreviations

ND: Non-Detectable

GaMD: Gaussian accelerated Molecular Dynamics

Lasso: least absolute shrinkage and selection operator

rRNA: ribosomal RNA

pre-mRNA: precursor messenger RNA

snRNP: small nuclear ribonucleoprotein

MD: molecular dynamics

lncRNA: long non-coding RNA

IRES: internal ribosome entry site

UTR: untranslated region of mRNA

HCV: hepatitis C virus

TAR: transactivation response

RNase L: ribonuclease L

RIBOTAC: ribonuclease targeting chimera

SAR: structure-activity relationship

FP: fluorescence polarization

dsDNA: double-stranded DNA

SOFAIR: band-Selective Optimized Flip-Angle Internally-encoded Relaxation

CPMG: Carr-Purcell-Meiboom-Gill

RMSFs: root-mean-square fluctuations

Fab: fragment antigen-binding region

QSAR: quantitative structure-activity-relationship

MOE: Molecular Operating Environment

DFT: density-functional theory

Lasso: least absolute shrinkage and selection operator

Ln$K_d$: natural logarithm of the binding constant

### Supplementary Figures

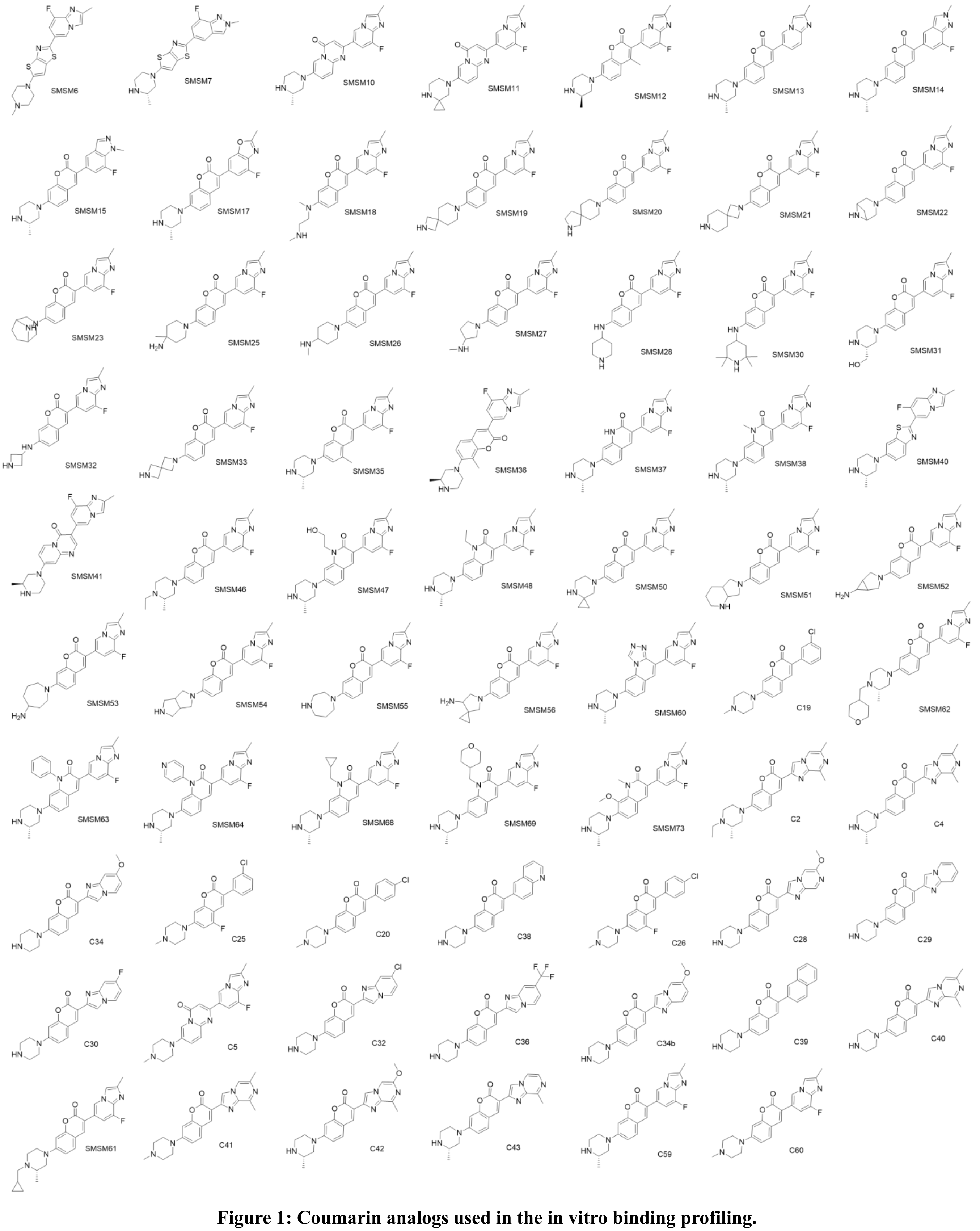

**Figure 1: Coumarin analogs used in the in vitro binding profiling.**

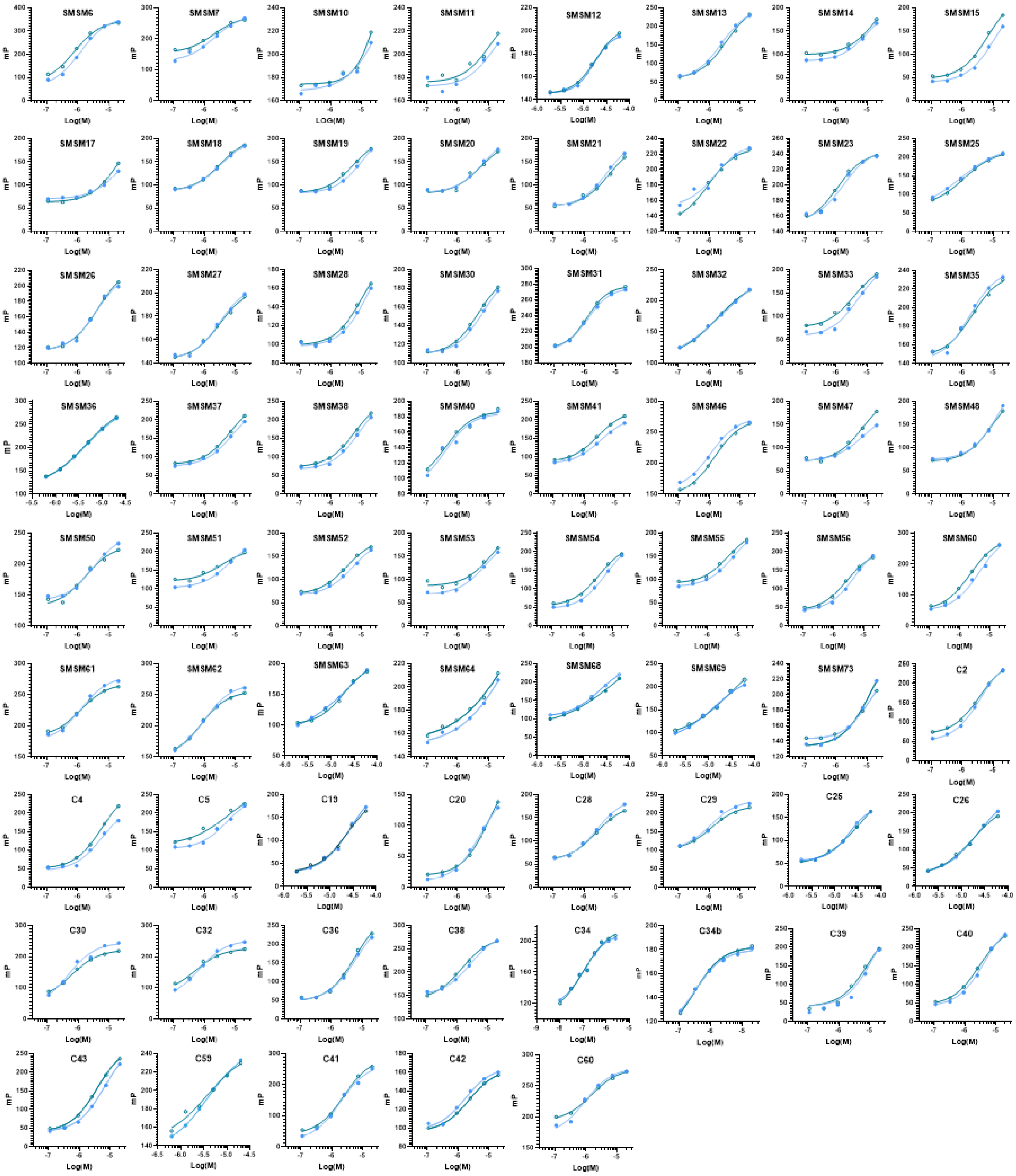

**Figure 2: Dose-response curves of in vitro binding of coumarin analogs and bulge G RNA (RNA1).** Two curves in each figure represent two independent experiments. Curve fitting was performed using a nonlinear regression function in GraphPad Prism 10.

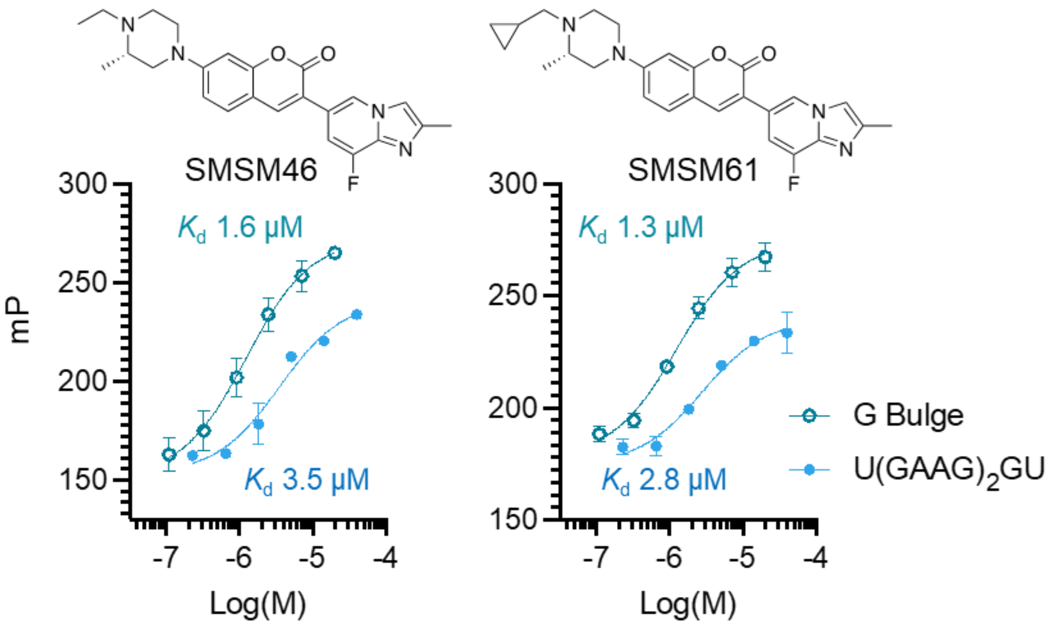

**Figure 3: Dose-response curves for compounds (SMSM46, SMSM61) selectively binding to bulged G RNA (RNA1) with a comparable binding affinity with an 11-nucleotide GA-rich sequence that would form a double loop-like RNA structure.**

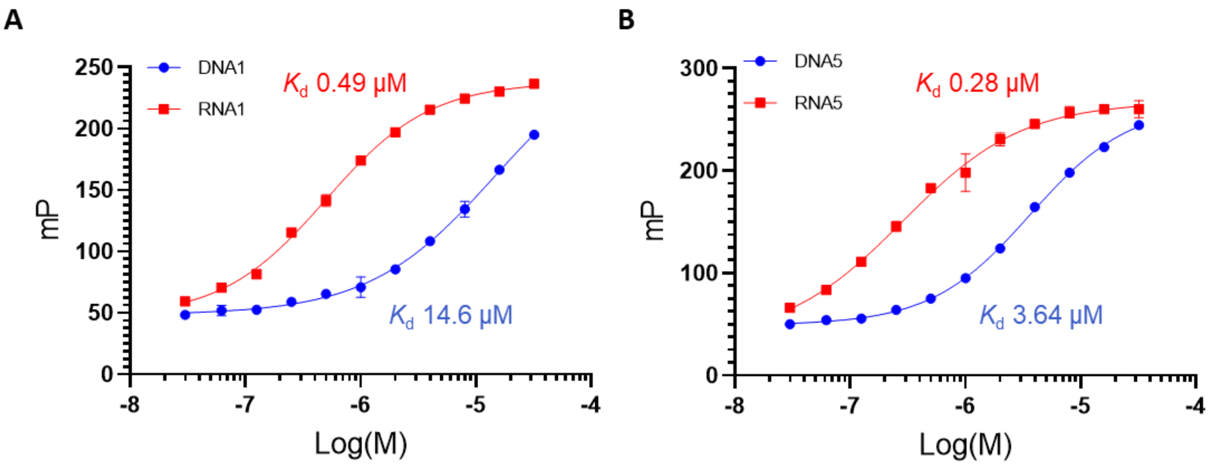

**Figure 4: Dose-response curves for C30 binding: (A)** Dose-response curves for C30 binding to RNA1 compared to its DNA counterpart sequence DNA1. **(B)** Dose-response curves for C30 binding to RNA5 compared to its DNA counterpart sequence DNA5.

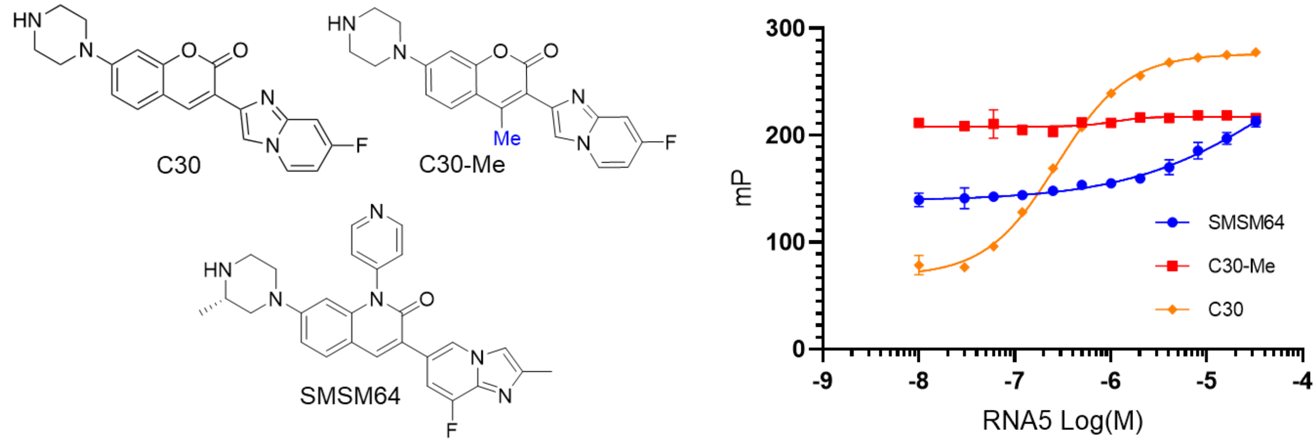

**Figure 5: Dose-response curves for C30, C30-Me and SMSM64 binding to RNA5**

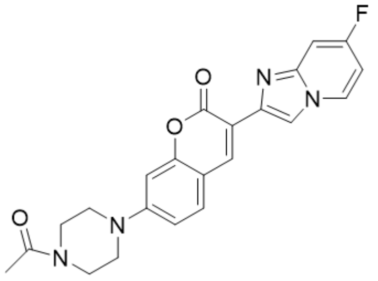

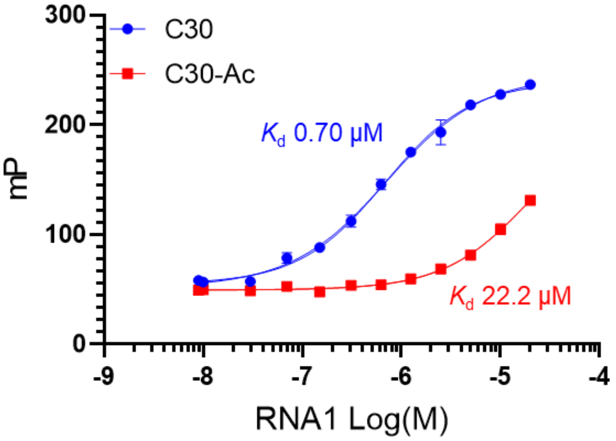

**Figure 6: Binding affinity of C30-Ac to RNA1:** (A) Structure of C30-Ac. (B) Dose-response curves for C30-Ac binding to RNA1

### Supplementary Tables

**Table 1: Observed binding affinity (Kd) of coumarin derivatives and RNA1 determined by fluorescence polarization.**

| Compound | $K_d$ (μM) | Compound | $K_d$ (μM) | Compound | $K_d$ (μM) |
|---|---|---|---|---|---|
| SMSM6 | 0.99±0.2 | C4 | 5.8±0.1 | SMSM48 | 11.8±2.3 |
| SMSM7 | 1.97±0.01 | C40 | 3.5±0.6 | SMSM50 | 2.4±0.6 |
| SMSM10 | 45.1±11.8 | C41 | 2.1±0.2 | SMSM51 | 4.9±1.3 |
| SMSM11 | 12.4±1.3 | C5 | 5.6±0.5 | SMSM52 | 4.1±0.9 |
| SMSM13 | 3.0±0.5 | C60 | 1.2±0.2 | SMSM53 | 10.0±2.1 |
| SMSM14 | 13.6±2.9 | C29 | 1.1±0.1 | SMSM54 | 4.6±1.1 |
| SMSM15 | 8.6±2.1 | C30 | 0.54±0.07 | SMSM55 | 5.9±1.0 |
| SMSM17 | 19.0±0.1 | C32 | 0.59±0.07 | SMSM56 | 3.9±1.1 |
| SMSM18 | 3.1±0.1 | C34 | 0.81±0.03 | SMSM60 | 3.3±1.1 |
| SMSM19 | 8.0±2.7 | C36 | 5.8±0.2 | C34b | 0.32±0.07 |
| SMSM20 | 5.9±0.5 | C43 | 5.0±1.3 | SMSM61 | 1.2±0.1 |
| SMSM21 | 5.9±0.4 | C42 | 1.9±0.2 | SMSM62 | 0.86±0.08 |
| SMSM22 | 1.2±0.3 | C28 | 2.4±0.1 | SMSM64 | 49.8±2.1 |
| SMSM23 | 1.3±0.2 | C38 | 2.0±0.6 | SMSM73 | 11.9±1.0 |
| SMSM25 | 1.5±0.5 | C39 | 10.5±2.1 | SMSM12 | 17.3±0.5 |
| SMSM26 | 4.0±0.2 | C20 | 7.9±2.5 | C19 | 25.3±1.4 |
| SMSM27 | 3.0±0.1 | SMSM35 | 1.9±0.1 | C25 | 27.9±7.5 |
| SMSM28 | 12.4±2.5 | SMSM37 | 7.1±0.2 | C26 | 19.7±2.0 |
| SMSM30 | 5.6±1.0 | SMSM38 | 6.9±1.0 | SMSM63 | 23.6±4.8 |
| SMSM31 | 1.2±0.3 | SMSM40 | 0.40±0.01 | SMSM68 | 30.5±7.8 |
| SMSM32 | 2.6±0.7 | SMSM41 | 3.0±0.2 | SMSM69 | 20.8±7.6 |
| SMSM33 | 4.4±0.6 | SMSM46 | 1.2±0.1 | C59 | 3.8±0.1 |
| C2 | 3.6±0.1 | SMSM47 | 6.9±0.1 | SMSM36 | 4.5±0.1 |

**Table 2: Crystallographic data for the representative Fab-apo RNA complex.**

| | Fab-RNA |
|---|---|
| Data Collection | |
| Unit-cell parameters (Å, °) | *a*=64.73, *b*=68.03, *c*=118.61, □=100.15 |
| Space group | $P2_1$ |
| Resolution (Å)[a] | 47.41-1.90 (1.94-1.90) |
| Wavelength (Å) | 0.9785 |
| Temperature (K) | 100 |
| Observed reflections | 364,335 |
| Unique reflections | 79,997 |
| <I/□(I)> [a] | 7.7 (1.7) |
| Completeness (%)[a] | 99.9 (100) |
| Multiplicity[a] | 4.6 (4.7) |
| $R_{merge}$ (%)[a,b] | 15.2 (108.9) |
| $R_{meas}$ (%)[a,d] | 17.1 (122.4) |
| $R_{pim}$ (%)[a,d] | 7.7 (55.0) |
| $CC_{1/2}$ [a,e] | 0.996 (0.625) |
| Refinement | |
| Resolution (Å) [a] | 47.41-1.90 |
| Reflections (working/test) [a] | 75,951/3,979 |
| $R_{factor}$ / $R_{free}$ (%)[a,c] | 17.6/22.3 |
| No. of atoms (Protein/RNA/Water) | 6,503/860/593 |
| Model Quality | |
| R.m.s deviations | |
| Bond lengths (Å) | 0.009 |
| Bond angles (°) | 0.899 |
| Mean *B*-factor (Å$^2$) | |
| All Atoms | 30.9 |
| Protein | 28.0 |
| RNA | 49.7 |
| Water | 33.1 |
| Coordinate error (maximum likelihood) (Å) | 0.21 |
| Ramachandran Plot | |
| Most favored (%) | 97.9 |
| Additionally allowed (%) | 2.1 |

[a] Values in parenthesis are for the highest resolution shell.
[b] $R_{merge}$ = □$_{hkl}$□$_i$ |$I_i(hkl)$ - <$I(hkl)$>| / □$_{hkl}$□$_i$ $I_i(hkl)$, where $I_i(hkl)$ is the intensity
[c] measured for the *i*th reflection and <$I(hkl)$> is the average intensity of all reflections with indices hkl.
[d] $R_{factor}$ = □$_{hkl}$ ||$F_{obs}$ $(hkl)$| - |$F_{calc}$ $(hkl)$|| / □$_{hkl}$ |$F_{obs}$ $(hkl)$|; Rfree is calculated in an
[e] identical manner using 5% of randomly selected reflections that were not included in the refinement.
[f] $R_{meas}$ = redundancy-independent (multiplicity-weighted) $R_{merge}$.[1,2] $R_{pim}$ = precision-indicating (multiplicity-weighted) $R_{merge}$.[3,4]
[g] $CC_{1/2}$ is the correlation coefficient of the mean intensities between two random half-sets of data.[5,6]

**Table 3: RNA5 1H peak assignment**

| RNA seq | | | chemical shift (ppm) |
|---|---|---|---|
| 22 | U | NH | 14.167 |
| 29 | U | NH | 13.893 |
| 5 | U | NH | 13.424 |
| 9 | G | NH | 13.194 |
| 25 | U | NH | 13.748 |
| 13 | G | NH | 12.483 |
| 24 | G | NH | 12.258 |
| 3 | G | NH | 12.176 |
| 11 | G | NH | 12.176 |
| 7 | G | NH | 11.654 |
| 27 | U | NH | 11.418 |
| 6 | G | NH | 10.117 |
| 10 | A | CH | 7.271 |
| 23 | C | NH2 | 6.849 |
| 21 | C | NH2 | 6.801 |
| 30 | C | NH2 | 6.699 |
| 26 | C | NH2 | 6.661 |

#### Table 4: RNA 1H R2 measurements (s-1) at varied C30 ligand concentrations

| | $R_2$ measured on RNA $^1$H resonances ($s^{-1}$) | | | | | | | | | | | | | |
|---|---|---|---|---|---|---|---|---|---|---|---|---|---|---|
| **NMR sample** | G3-NH | U5-NH | G6-NH | G7-NH | G9-NH | A10-CH | C21-$NH_2$ | U22-NH | C23-$NH_2$ | G24-NH | C26-$NH_2$ | U27-NH | U29-NH | C30-$NH_2$ |
| 700 $\mu$M RNA | 115.98 | 62.33 | 119.74 | — | 45.01 | 18.69 | 76.71 | 51.78 | 32.45 | 72.05 | 82.18 | 101.02 | 49.80 | 101.46 |
| 700 $\mu$M RNA + 35 $\mu$M ligand | 114.16 | 63.50 | 115.16 | — | 50.73 | 22.52 | 75.83 | 55.69 | 36.71 | 74.46 | 82.76 | 96.94 | 49.82 | 100.57 |
| 700 $\mu$M RNA + 70 $\mu$M ligand | 113.46 | 63.94 | 116.39 | 108.39 | 55.25 | 25.44 | 79.83 | 58.68 | 44.33 | 76.89 | 88.06 | 94.84 | 51.74 | 105.06 |
| 700 $\mu$M RNA + 140 $\mu$M ligand | 112.55 | 64.82 | 115.10 | 108.80 | 63.04 | 30.18 | 83.27 | 63.64 | 53.91 | 82.12 | 91.51 | 91.99 | 51.99 | 106.18 |
| 700 $\mu$M RNA + 280 $\mu$M ligand | 107.28 | 67.55 | 113.47 | 107.80 | 80.18 | 42.03 | 89.82 | 75.48 | 80.34 | 92.79 | 97.37 | 90.28 | 55.72 | 109.12 |
| 700 $\mu$M RNA + 350 $\mu$M ligand | 107.55 | 70.08 | 114.12 | 105.64 | 92.72 | 50.69 | 97.69 | 83.14 | 98.19 | 103.14 | 102.87 | 89.42 | 57.16 | 113.25 |
| 700 $\mu$M RNA + 490 $\mu$M ligand | 106.60 | 71.76 | 115.84 | 106.54 | 111.64 | 63.67 | 96.74 | 101.06 | 109.98 | 122.20 | 103.46 | 88.17 | 57.97 | 113.38 |
| 700 $\mu$M RNA + 700 $\mu$M ligand | 100.32 | 76.14 | 117.15 | 120.62 | 155.42 | 73.75 | 101.06 | 135.92 | 121.97 | 147.85 | 105.25 | 88.91 | 60.59 | 118.33 |

**Table 5: Non-zero coefficients for LnKd determined through Lasso regression.**

| Molecular descriptor | Class | Coefficient | Description[a] | Impact[b] |
|---|---|---|---|---|
| FCharge | charge | -0.004721127 | Total charge of the molecule (sum of formal charges). | + |
| a_base | charge | -0.253475222 | Number of basic atoms. | + |
| PEOE_VSA_FPOS | charge | -0.0001459042 | Fractional positive VDWSA. | + |
| PEOE_VSA_FNEG[c] | charge | 1.9606574824 | Fractional negative VDWSA. | − |
| PEOE_VSA_NEG | charge | 0.0027418015 | Total negative VDWSA, i.e., sum of the $v_i$ such that $q_i$ is negative, where $v_i$ = the accessible VDWSA (in Å$^2$) and $q_i$ = the partial charge calculation for each atom. | − |
| BCUT_SMR_1 | charge | -1.542968189 | The BCUT descriptors are calculated from the eigenvalues of a modified adjacency matrix. Each *ij* entry of the adjacency matrix takes the value 1/sqrt($b_{ij}$) where $b_{ij}$ is the formal bond order between bonded atoms *i* and *j*. The diagonal takes the value of the PEOE partial charges. The resulting eigenvalues are sorted and the smallest, 1/3-ile, 2/3-ile and largest eigenvalues are reported. The BCUT descriptors are calculated using atomic contribution to molar refractivity instead of partial charge. | + |
| NPR1 | shape | 2.9102299092 | Normalized principal moments of inertia (PMI) ratio: PMI1/PMI3. | − |
| std_dim1 | shape | -0.2822878479 | Standard dimension 1: the square root of the largest eigenvalue of the covariance matrix of the atomic coordinates. A standard dimension is equivalent to the standard deviation along a principal component axis. | + |
| $\omega$(BC-DE) | shape | 0.0079825491 | Dihedral in the optimal structure between BC and DE rings. | – |
| E_str | energy | 0.0660924541 | Bond stretch potential energy. | – |
| PM3_HOMO | energy | 0.0027418015 | The energy (eV) of HOMO calculated using the PM3 Hamiltonian. | – |
| vsurf_EDmin3 | energy | -0.994318259 | Vsurf_EDmin describes the lowest hydrophobic energy representing the distances between the best three local minima of interaction energy when a hydrophobic probe (DRY) interacts with the target molecule. | + |
| MNDO_dipole | other | 0.0020593997 | The dipole moment calculated using the MNDO Hamiltonian. | – |
| SlogP_VSA0 | other | -3.021185e-05 | Sum of $v_i$ such that $L_i$ <= -0.4, where $L_i$ = the contribution to logP(o/w) for atom *i*. | + |
| SlogP_VSA6 | other | 0.1114724012 | Sum of $v_i$ such that $L_i$ is in (0.20,0.25]. | – |
| SMR_VSA6 | other | -0.002865974 | Sum of $v_i$ such that $R_i$ is in (0.485,0.56]. $R_i$ = the contribution to Molar Refractivity for atom *i*. | + |

[a] VDWSA = van der Waals surface area. [b] + and – signs indicate variables positively or negatively correlated to the binding affinity. [c] PEOE_VSA_FNEG = – PEOE_VSA_FPOS

### Methods

#### DFT Calculations

The DFT calculations were performed using the Gaussian 09 software (Revision C.01).

Reference for Gaussian 09 software.
M. J. Frisch, G. W. Trucks, H. B. Schlegel, G. E. Scuseria, M. A. Robb, J. R. Cheeseman, G. Scalmani, V. Barone, B. Mennucci, G. A. Petersson, H. Nakatsuji, M. Caricato, X. Li, H. P. Hratchian, A. F. Izmaylov, J. Bloino, G. Zheng, J. L. Sonnenberg, M. Hada, M. Ehara, K. Toyota, R. Fukuda, J. Hasegawa, M. Ishida, T. Nakajima, Y. Honda, O. Kitao, H. Nakai, T. Vreven, J. A. Montgomery, Jr., J. E. Peralta, F. Ogliaro, M. Bearpark, J. J. Heyd, E. Brothers, K. N. Kudin, V. N. Staroverov, T. Keith, R. Kobayashi, J. Normand, K. Raghavachari, A. Rendell, J. C. Burant, S. S. Iyengar, J. Tomasi, M. Cossi, N. Rega, J. M. Millam, M. Klene, J. E. Knox, J. B. Cross, V. Bakken, C. Adamo, J. Jaramillo, R. Gomperts, R. E. Stratmann, O. Yazyev, A. J. Austin, R. Cammi, C. Pomelli, J. W. Ochterski, R. L. Martin, K. Morokuma, V. G. Zakrzewski, G. A. Voth, P. Salvador, J. J. Dannenberg, S. Dapprich, A. D. Daniels, O. Farkas, J. B. Foresman, J. V. Ortiz, J. Cioslowski, and D. J. Fox, Gaussian, Inc., Wallingford CT, 2010.

#### Crystallization and Data Collection

Purified Fab protein was complexed with RNA (5' CUCGUCUGAAACACGGAGAG 3') at a concentration of 2.5 mg/mL for crystallization screening. All crystallization experiments were set up using an NT8 drop-setting robot (Formulatrix) and UVXPO MRC (Molecular Dimensions) sitting drop vapor diffusion plates at 18 °C. 100 nL of protein and 100 nL crystallization solution were dispensed and equilibrated against 50 uL of the latter. Crystals were obtained in 1-2 days from the Index HT screen (Hampton Research) condition A6 (2M ammonium sulfate, 100 mM Tris pH 8.5). Samples were transferred to a fresh drop composed of 80% crystallization solution and 20% (v/v) glycerol and stored in liquid nitrogen. X-ray diffraction data were collected at the National Synchrotron Light Source II (NSLS-II) beamline 19-ID (NYX) using a Dectris Eiger2 × 9M pixel array detector. Mosquito liquid handling robot (SPT Labtech) was used to set up crystal trials of RNA structures aligned in S10 with the purified Fab. 100 nl + 100 nl hanging drop crystal trials were set in commercially available crystallization kits from Hampton Research and Jena Bioscience and allowed to grow for 2−3 weeks at room temperature. Multiple crystals grew in variety of conditions. Crystals were then looped and frozen in liquid nitrogen. X-ray diffraction data were collected at the National Synchrotron Light Source II (NSLS-II) beamline 17-ID (FMX/AMX).

#### Crystal Structure Solution and Refinement

Intensities were integrated using XDS [7,8] via Autoproc[9] and the Laue class analysis and data scaling were performed with Aimless.[1] Structure solution was conducted by molecular replacement with Phaser [10] using a previously determined structure of the heavy and light chains from a Fab-RNA structure (PDB 6B3K) as the search model. Structure refinement and manual model building were conducted with Phenix [11] and Coot [12] respectively. Disordered side chains were truncated to the point for which electron density could be observed. Structure validation was conducted with Molprobity [13] and figures were prepared using the CCP4MG package.[14]

### Chemistry

#### General Methods

Reagents and solvents were purchased from commercial sources (Fisher, Sigma-Aldrich and Combi-Blocks) and used as received. Reactions were tracked by TLC (Silica gel 60 $F_{254}$, Merck) and Waters ACQUITY UPLC-MS system (ACQUITY UPLC H Class Plus in tandem with Qda Mass Detector). Intermediates and products were purified by a Teledyne ISCO Combi-Flash system using prepacked $SiO_2$ cartridges. NMR spectra were acquired on a Bruker AV400 instrument (400 MHz for $^1$H NMR, 100 MHz for $^{13}$C NMR). $^{13}$C shifts were obtained with $^1$H decoupling. MestReNova 14.0.1 developed by MESTRELAB RESEARCH was used for NMR data processing. MS-ESI spectra were recorded on Waters Qda Mass Detector. HRMS-ESI were recorded on a Agilent 6224 mass spectrometer with a time-of-flight (TOF) analyzer.

Compounds **C30**, **C34, Int1, Int2 and S6** were synthesized following procedures reported in literature and verified by NMR and Mass spectra.[15,16]

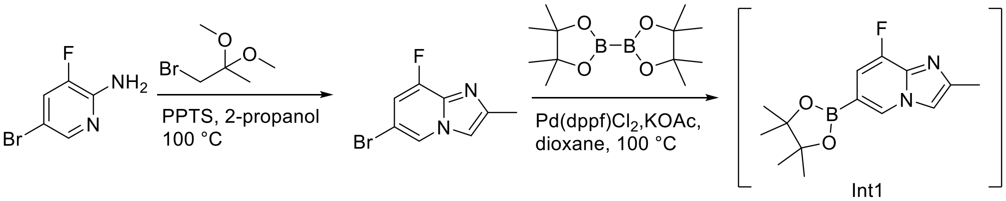

Int1

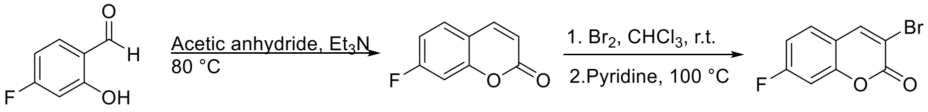

Int2

**2-(8-fluoro-2-methylimidazo[1,2-a]pyridin-6-yl)-5-(4-methylpiperazin-1-yl)thieno[2,3-d]thiazole (SMSM6)**

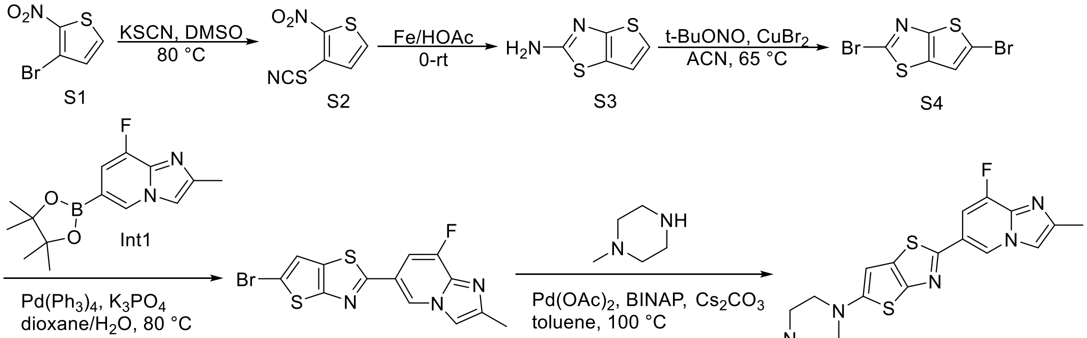

To a solution of S1 (1.4 g, 6.7 mmol) in DMSO (15 mL) was added potassium thiocyanate (1.96 g, 20 mmol). The mixture was heated at 80 °C for 4 h. After cooled to room temperature, the mixture was added to ice-water and extracted with ethyl acetate (EA). The organic layer was washed with brine and dried over sodium sulfate. EA was removed under vacuum and the resulting product was used directly in the next step. (1.2 g, yield 96%). MS-ESI (*m/z*) $[M+H]^+$ 187.02.

To a solution of S2 (1.2 g, 6.4 mmol) in acetic acid (20 mL) was added iron powder (1.8 g, 32 mmol) in portions. The mixture was stirred at room temperature overnight. Water was added and the mixture was filtered. The filtrate was adjusted to pH 8 using saturated aq$Na_2CO_3$ and extracted with EA. The organic layer was washed with brine and dried over sodium sulfate. EA was removed under vacuum and the crude product was purified by silica gel column chromatography (0 – 100% EA in hexanes) to give the pure product. (0.8, yield 80%). MS-ESI (*m/z*) $[M+H]^+$ 157.0.

To a solution of S3 (0.8 g, 5.1 mmol) in acetonitrile (15 mL) was added $CuBr_2$ (1.1 g, 5.1 mmol) and tert-butyl nitrite (0.9 mL, 7.6 mmol) dropwise. The mixture was heated at 65 °C for 1 h. after cooled to room temperature, the mixture was diluted with EA and filtered through Celite. The filtrate was washed with brine and dried over sodium sulfate. Solvent was removed under vacuum and the crude product was purified by silica gel column chromatography (0 – 10% EA in hexanes) to give the pure product. (0.2, yield 13%). MS-ESI (*m/z*) $[M+H]^+$ 299.8.

To a solution of S4 (0.2 g, 0.66 mmol) and 8-fluoro-2-methyl-6-(4,4,5,5-tetramethyl-1,3,2-dioxaborolan-2-yl)imidazo[1,2-a]pyridine (0.27 g, 1 mmol) in dioxane (5 mL) was added $Pd(PPh_3)_4$ (68 mg, 0.066 mmol), $K_2CO_3$ (0.21 g, 1.98 mmol) and water (1 mL). The mixture was degassed and refilled with nitrogen and then heated at 80 ℃ overnight. Ethyl acetate (10 mL) was added, the mixture was filtered through Celite. The organic layer was washed with brine, dried over sodium sulfate. Solvent was removed under vacuum and the crude product was purified by silica gel column chromatography (0 – 100% EA in hexanes) to give the pure product. (0.1 g, yield 41%). MS-ESI (*m/z*) $[M+H]^+$ 367.9.

A mixture of S5 (0.1 g, 0.27 mmol), N-methylpiperazine (60 µL, 0.54 mmol), $Pd(OAc)_2$ (6 mg, 0.027 mmol), BINAP (33 mg, 0.054 mmol) and $Cs_2CO_3$ (0.26 g, 0.81 mmol) in toluene (2 mL) was degassed and refilled with nitrogen and then heated at 80 ℃ overnight. Ethyl acetate (5 mL) was added, the mixture was filtered through Celite. Solvent was removed under vacuum and the crude product was purified by silica gel column chromatography (0 – 10% MeOH in DCM) to give SMSM6 as a yellow solid. (20 mg, yield 19%). HRMS-ESI (m/z) $[M+Na]^+$ Calcd 410.0885; Found 410.0857.

$^1$H NMR (400 MHz, DMSO-$d_6$) δ 9.03 (d, *J* = 1.5 Hz, 1H), 7.88 (d, *J* = 2.1 Hz, 1H), 7.56 (dd, *J* = 11.9, 1.5 Hz, 1H), 6.52 (s, 1H), 3.19 (t, *J* = 4 Hz, 4H), 2.48 (t, *J* = 4 Hz, 4H), 2.37 (s, 3H), 2.24 (s, 3H).

$^{13}$C NMR (100 MHz, DMSO-$d_6$) δ 160.2, 159.4 (d, *J* = 2.8 Hz), 150.2 (d, *J* = 249.4 Hz), 145.1, 144.2, 136.4 (d, *J* = 28.3 Hz), 132.5, 120.8 (d, *J* = 4.3 Hz), 119.2 (d, *J* = 7.3 Hz), 113.4, 105.3 (d, *J* = 18.5 Hz), 95.2, 54.2, 50.6, 46.1, 14.7.

**(*S*)-7-(4-ethyl-3-methylpiperazin-1-yl)-3-(8-fluoro-2-methylimidazo[1,2-*a*]pyridin-6-yl)-2*H*-chromen-2-one (SMSM46)**

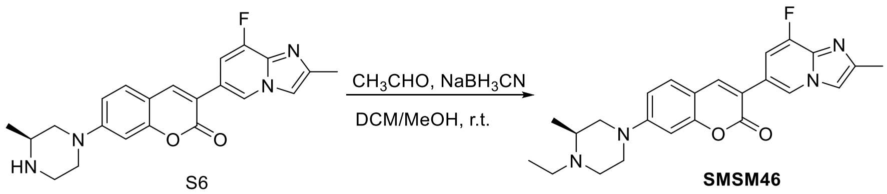

To a solution of S6 (10 mg, 0.025 mmol) in DCM/MeOH (0.2/0.2 mL) was added acetaldehyde (4.2 µL, 0.075 mmol) and $NaBH_3CN$ (4.7 mg, 0.075 mmol). The mixture was stirred at room temperature for 2 h. Water was added, and the mixture was extracted with DCM. The crude product was purified by silica gel column chromatography (0 – 10% MeOH in DCM) to give **SMSM46** as a yellow solid. (4 mg, yield 38%). HRMS-ESI (m/z) $[M+H]^+$ Calcd 421.2040; Found 421.2032.

$^1$H NMR (400 MHz, DMSO-$d_6$) δ 8.91 (d, *J* = 1.4 Hz, 1H), 8.29 (s, 1H), 7.90 (dd, *J* = 3.2, 1.1 Hz, 1H), 7.54 (d, *J* = 8.9 Hz, 1H), 7.48 (dd, *J* = 13.0, 1.5 Hz, 1H), 7.02 (dd, *J* = 8.9, 2.4 Hz, 1H), 6.89 (d, *J* = 2.3 Hz, 1H), 3.75 – 3.70 (m, 2H), 3.07 – 3.00 (m, 1H), 2.89 – 2.84 (m, 1H), 2.81 – 2.70 (m, 2H), 2.45 (brs, 1H), 2.36 (s, 3H), 2.34 – 2.26 (m, 2H), 1.06 (d, *J* = 6.2 Hz, 3H), 0.98 (t, *J* = 7.1 Hz, 3H).

$^{13}$C NMR (100 MHz, DMSO-$d_6$) δ 160.5, 155.7, 153.8, 149.8 (d, *J* = 247.2 Hz), 143.6, 141.5, 135.9 (d, *J* = 28.9 Hz), 129.9, 122.5, 119.2, 116.3, 113.1, 112.0, 110.4, 107.6 (d, *J* = 17.4 Hz), 99.5, 54.1, 53.9, 49.6, 47.3, 46.8, 16.0, 14.7, 11.1.

**(*S*)-7-(4-(cyclopropylmethyl)-3-methylpiperazin-1-yl)-3-(8-fluoro-2-methylimidazo[1,2-*a*]pyridin-6-yl)-2*H*-chromen-2-one (SMSM61)**

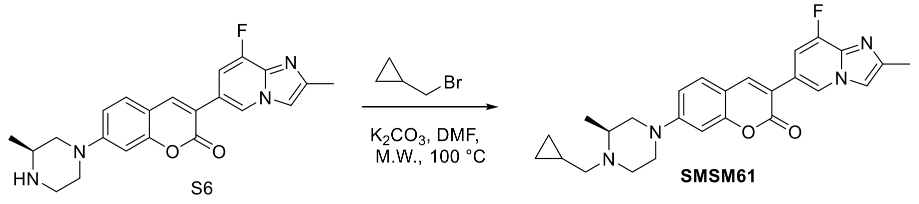

To a solution of S6 (5 mg, 0.0125 mmol) in DMF (0.2 mL) was added (bromomethyl)cyclopropane (2.5 μL, 0.025 mmol) and $K_2CO_3$ (4 mg, 0.0375 mmol). The mixture was heated at 100 °C for 1 h in a Biotage Initiator+ microwave reactor. DMF was removed under vacuum, the residue was purified by silica gel column chromatography (0 – 10% MeOH in DCM) to give **SMSM61** as a yellow solid. (4 mg, yield 71%). HRMS-ESI (m/z) $[M+H]^+$ Calcd 447.2196; Found 447.2181.

$^1$H NMR (400 MHz, DMSO-$d_6$) δ 8.92 (d, *J* = 1.4 Hz, 1H), 8.29 (s, 1H), 7.90 (dd, *J* = 3.1, 1.1 Hz, 1H), 7.54 (d, *J* = 8.9 Hz, 1H), 7.48 (dd, *J* = 12.9, 1.5 Hz, 1H), 7.03 (dd, *J* = 8.9, 2.4 Hz, 1H), 6.90 (d, *J* = 2.3 Hz, 1H), 3.75 (t, *J* = 12.3 Hz, 2H), 3.10 – 3.03 (m, 2H), 2.77 – 2.71 (m, 1H), 2.61 (dd, *J* = 13.1, 6.4 Hz, 1H), 2.42 – 2.36 (d, *J* = 0.8 Hz, 4H), 2.20 – 2.12 (m, 1H), 1.06 (d, *J* = 6.1 Hz, 3H), 0.87 – 0.81 (m, 1H), 0.53 – 0.42 (m, 2H), 0.14 – 0.06 (m, 2H).

$^{13}$C NMR (100 MHz, DMSO-$d_6$) δ 160.5, 155.7, 153.8, 149.8 (d, *J* = 246.9 Hz), 143.6, 141.5, 135.9 (d, *J* = 28.9 Hz), 129.9, 122.6, 119.2 (d, *J* = 7.6 Hz), 116.3, 113.1, 112.0, 110.4, 107.6 (d, *J* = 17.5 Hz), 99.5, 57.9, 54.6, 53.9, 50.6, 47.3, 16.1, 14.7, 8.0, 4.9, 3.6.

**(*S*)-3-(8-fluoro-2-methylimidazo[1,2-*a*]pyridin-6-yl)-7-(3-methylpiperazin-1-yl)-1-(pyridin-4-yl)quinolin-2(1*H*)-one (SMSM64)**

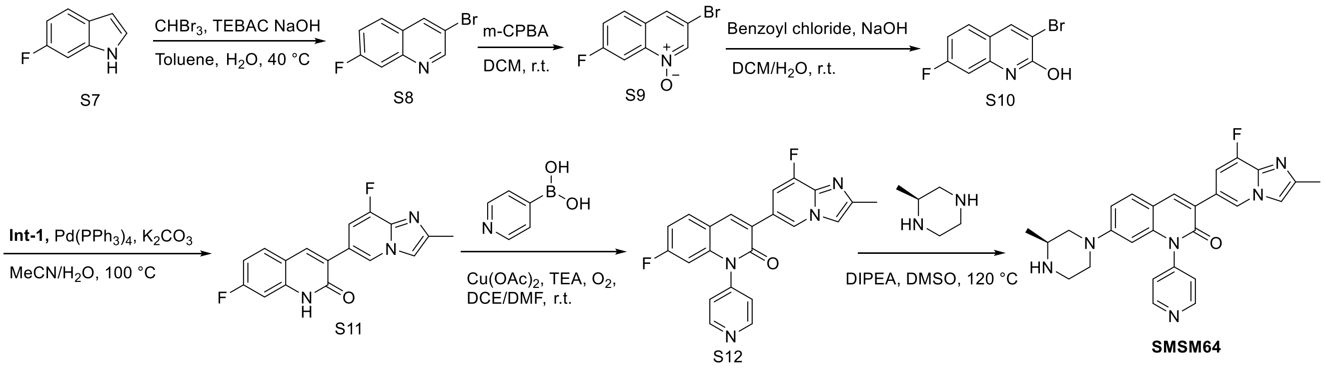

To a stirred mixture of S7 (4.9 g, 36 mmol), bromoform (3.2 mL, 36 mmol), and TEBAC (0.42 g, 1.8 mmol) in toluene (4 mL) at 40°C was added sodium hydroxide (11.6 g, 290 mmol) in water (22 mL) slowly, the resulting mixture was stirred at 40 °C overnight. The reaction mixture was cooled to room temperature, poured into water (100 mL), and extracted with DCM. The combined organic phase was washed with brine and dried over anhydrous sodium sulfate. The organic phase was concentrated under vacuum and the crude product was purified by silica gel flash column chromatography using 3% EA in hexanes to give S8 (1.45 g, yield 18%) as a white solid. MS-ESI (*m*/*z*) $[M+H]^+$ 226.05.

$^1$H NMR (400 MHz, $CDCl_3$): δ 8.91 (d, *J* = 2.0 Hz, 1H), 8.31 (d, *J* = 2.4 Hz, 1H), 7.77 – 7.70 (m, 2H), 7.40 – 7.35 (m, 1H).

To a solution of S8 (1 g, 4.4 mmol) in DCM (25 mL) was added *m*-CPBA (0.36 g, 1.77 mmol). The mixture was stirred at room temperature overnight. Water was added and the mixture was extracted with DCM. The organic layer was washed with brine, dried over anhydrous sodium sulfate, and concentrated. The crude product was purified by silica gel column chromatography (0 – 25% EA in hexanes) to give the pure product as a light yellow solid (0.8 g, yield 75%). MS-ESI (*m*/*z*) $[M+H]^+$ 242.0.

$^1$H NMR (400 MHz, $CDCl_3$): δ 8.63 (d, *J* = 0.8 Hz, 1H), 8.35 – 8.32 (m, 1H), 7.90 (s, 1H), 7.84 – 7.81 (m, 1H), 7.48 – 7.43 (m, 1H).

To a stirred mixture of S9 (0.8 g, 3.36 mmol) and sodium hydroxide (0.32 g, 7.76 mmol) in water (5 mL) and DCM (10 mL) at room temperature was added benzoyl chloride (0.56 g, 4 mmol) dropwise. The resulting mixture was stirred at room temperature for 2 h. The reaction mixture was filtered, washed with water and dried to give S10 as a white solid (0.43 g, yield 54%). MS-ESI (*m*/*z*) $[M+H]^+$ 242.05.

$^1$H NMR (400 MHz, DMSO-$_{d6}$): δ 12.35 (s, 1H), 8.52 (s, 1H), 7.77 – 7.73 (m, 1H), 7.13 – 7.05 (m, 2H).

A mixture of S10 (0.15 g, 0.63 mmol), Int-1 (0.34 g, 1.2 mmol), $K_2CO_3$ (0.26 g, 1.89 mmol), and tetrakis(triphenylphosphine)palladium (72 mg, 0.06 mmol) in acetonitrile (5 mL) and water (1 mL) was stirred at 80 °C under nitrogen overnight. Water (4 mL) was added, and the mixture was filtered. The crude product was purified by silica gel column chromatography (0 – 5% MeOH in DCM) to give S11 as a white solid. (60 mg, yield 32%). MS-ESI (*m*/*z*) $[M+H]^+$ 312.5.

$^1$H NMR (400 MHz, DMSO-$_{d6}$): δ 12.20 (s, 1H), 9.08 (s, 1H), 8.36 (s, 1H), 7.92 (d, *J* = 2.4 Hz, 1H), 7.83 – 7.79 (m, 1H), 7.55 – 7.51 (m, 1H), 7.15 – 7.09 (m, 2H), 2.37 (s, 3H).

A mixture of S11 (60 mg, 0.19 mmol), 4-pyridinylboronic acid (70 mg, 0.57 mmol), $Cu(OAc)_2$ (68 mg, 0.38 mmol) and TEA (79 µL, 0.57 mmol) in dichloroethane (2 mL) and DMF (3 mL) was stirred at room temperature for 48 h under oxygen. The mixture was filtered and washed with DCM and MeOH. Solvent was removed under vacuum and the residue was purified by silica gel column chromatography (0 – 5% MeOH in DCM) to give S12 as a white solid. (8 mg, yield 11%). MS-ESI (*m*/*z*) $[M+H]^+$ 389.2.

A mixture of S12 (8 mg, 0.02 mmol), (*S*)-2-methylpiperazine (4 mg. 0.04 mmol) and $K_2CO_3$ (8 mg, 0.06 mmol) in DMSO (0.4 mL) was heated at 120 °C overnight. The mixture was added to ice-water and filtered. The crude product was purified by silica gel column chromatography (0 – 20% MeOH in DCM) to afford **SMSM64** as a yellow solid (4 mg, yield 43%). HRMS-ESI (*m*/*z*) $[M+H]^+$ Calcd 469.2152; Found 469.2147.

$^1$H NMR (400 MHz, DMSO-$d_6$) δ 9.02 (d, *J* = 1.4 Hz, 1H), 8.88 (d, *J* = 6.0 Hz, 2H), 8.37 (s, 1H), 7.86 (d, *J* = 3.1 Hz, 1H), 7.71 (d, *J* = 8.9 Hz, 1H), 7.59 – 7.48 (m, 3H), 7.07 (dd, *J* = 8.9, 2.3 Hz, 1H), 5.78 (d, *J* = 2.2 Hz, 1H), 3.59 – 3.56 (m, 1H), 3.43 – 3.40 (m, 2H), 3.06 – 3.03 (m, 1H), 2.93 (brs, 1H), 2.82 – 2.71 (m, 2H), 2.35 (s, 3H), 1.08 (d, *J* = 6.3 Hz, 3H).

$^{13}$C NMR (100 MHz, DMSO-$d_6$) δ 160.1, 152.1, 151.9, 149.2 (d, *J* = 247.0 Hz), 145.9, 143.0, 138.0, 130.2, 124.5, 122.3, 121.1, 119.7, 112.5, 112.3, 111.7, 107.5 (d, *J* = 17.3 Hz), 98.4, 54.9, 52.6, 49.8, 45.5, 43.4, 14.7.

**7-(4-acetylpiperazin-1-yl)-3-(7-fluoroimidazo[1,2-*a*]pyridin-2-yl)-2*H*-chromen-2-one (C30-Ac)**

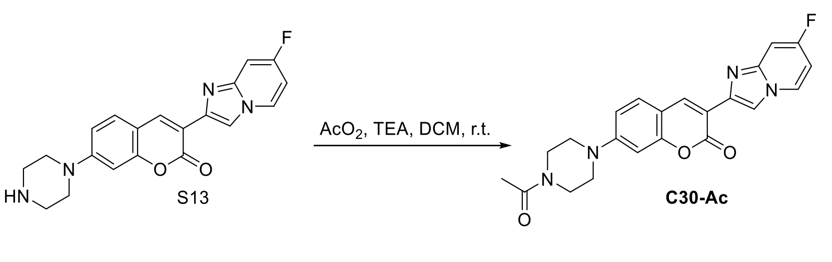

To a solution of C30 (6 mg, 0.016 mmol) in DCM (0.2 mL) was added acetic anhydride (4 µL, 0.032 mmol) and TEA (6.6 µL, 0.048 mmol). The mixture was stirred at room temperature for 2 h. DCM was removed, and the residue was purified by silica gel column chromatography (0 – 10% $CH_3OH$ in $CH_2Cl_2$) to give **C30-Ac** as a yellow solid. (5 mg, yield 77%). HRMS-ESI (*m*/*z*) $[M+H]^+$ Calcd 407.1519; Found 407.1513.

$^1$H NMR (400 MHz, DMSO-$d_6$) δ 8.71 – 8.68 (m, 2H), 8.52 (s, 1H), 7.72 (d, *J* = 8.9 Hz, 1H), 7.39 (dd, *J* = 10.1, 2.6 Hz, 1H), 7.04 (dd, *J* = 8.9, 2.4 Hz, 1H), 6.98 (td, *J* = 7.6, 2.6 Hz, 1H), 6.90 (d, *J* = 2.4 Hz, 1H), 3.61 – 3.59 (m, 4H), 3.48 – 3.45 (m, 2H), 3.41 – 3.38 (m, 2H), 2.06 (s, 3H).

$^{13}$C NMR (100 MHz, DMSO-$d_6$) δ 168.5, 160.2 (d, *J* = 250.0 Hz), 159.5, 154.9, 153.0, 144.4, 139.3, 138.6, 129.7, 129.3, 114.6, 112.0 (d, *J* = 29.9 Hz), 110.4, 104.2 (d, *J* = 30.1 Hz), 99.7, 99.6, 99.4, 46.8, 46.5, 45.0, 21.2.

#### 3-(7-fluoroimidazo[1,2-*a*]pyridin-2-yl)-5-methyl-7-(piperazin-1-yl)-2*H*-chromen-2-one (C30-Me$^{\text{Ring B}}$)

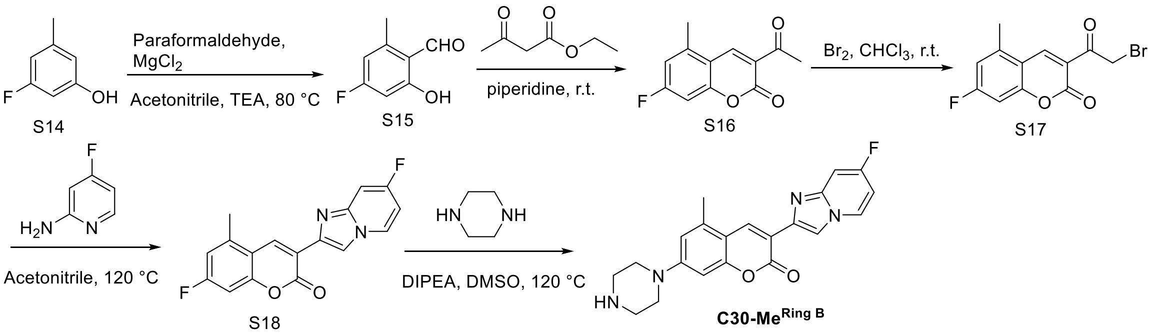

To a solution of 3-fluoro-5-methylphenol (1 g, 7.9 mmol) in acetonitrile (30 mL) was added paraformaldehyde (1.54 g, 51 mmol), $MgCl_2$ (1.1 g, 11.8 mmol), and TEA (2.1 mL, 29.6 mmol) respectively. The resulting mixture was stirred at 80 °C for 3 h. The reaction mixture was quenched with 2N HCl and extracted using ethyl acetate (EA). The crude product was purified by silica gel column chromatography (0-50% EA in hexanes) to afford S15 as a light yellow solid (0.4 g, yield 33%). MS-ESI (*m*/*z*) $[M+H]^+$ 155.0.

To S15 (0.4 g, 2.6 mmol) was added ethyl acetoacetate (0.33 mL, 2.6 mmol) and piperidine (25 µL, 0.26 mmol). The mixture was stirred at room temperature for 1 h. Precipitate formed during reaction. Methanol was added and the mixture was sonicated and filtered to give S16 as a yellow solid which was used without further purification (0.3 g, yield 52%). MS-ESI (*m*/*z*) $[M+H]^+$ 221.1.

To a solution of S16 (0.3 g, 1.36 mmol) in $CHCl_3$ (10 mL) was added bromine (77 µL, 1.5 mmol). The mixture was stirred at room temperature for 1 h. A colorless mixture was obtained. $CHCl_3$ was removed under vacuum and methanol was added. The suspension was sonicated and filtered to give S17 as a yellow solid (0.2 g, yield 49%). MS-ESI (*m*/*z*) $[M+H]^+$ 299.0.

Compound **C30-Me$^{\text{Ring B}}$** was synthesized following the same procedure for C34b by using S17 and 4-fluoro-2-aminopyridine. Yellow solid, 10 mg, two-step yield 30%. HRMS-ESI (*m*/*z*) $[M+H]^+$ Calcd 379.1570; Found 379.1559.

$^1$H NMR (400 MHz, DMSO-$d_6$) δ 8.69 – 8.66 (m, 2H), 8.50 (s, 1H), 7.42 (dd, *J* = 10.2, 2.6 Hz, 1H), 6.95 (td, *J* = 7.6, 2.6 Hz, 1H), 6.90 (d, *J* = 2.4 Hz, 1H), 6.69 (d, *J* = 2.4 Hz, 1H), 3.34 (t, *J* = 4.0 Hz, 4H), 2.88 (t, *J* = 4.0 Hz, 4H), 2.53 (s, 3H).

$^{13}$C NMR (100 MHz, DMSO-$d_6$) δ 160.6 (d, *J* = 247.0 Hz), 159.9, 156.0, 153.5, 144.8 (d, *J* = 14.5 Hz), 140.0, 137.8, 135.8, 129.6 (d, *J* = 11.3 Hz), 114.1, 113.2, 112.5, 109.7, 104.6 (d, *J* = 29.5 Hz), 100.1 (d, *J* = 23.8 Hz), 98.0, 47.6, 45.3, 19.0.

**3-(7-fluoroimidazo[1,2-*a*]pyridin-2-yl)-4-methyl-7-(piperazin-1-yl)-2*H*-chromen-2-one (C30-Me)**

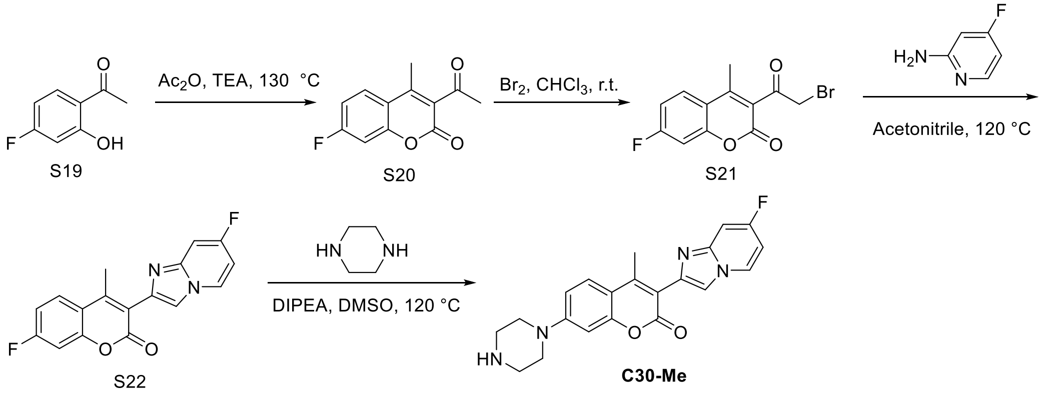

To 1-(4-fluoro-2-hydroxyphenyl)ethan-1-one (1 g, 6.5 mmol) was added acetic anhydride (3 mL) and triethylamine (1 mL). The mixture was heated at 130 ℃ for 48 h. The mixture was cooled to room temperature, poured into ice water. The sticky solid was isolated and dissolved in ethyl acetate. The organic layer was washed with brine, dried with anhydrous sodium sulfate and removed under vacuum. The crude product was purified by column chromatography (0 - 50% ethyl acetate in hexanes) to afford the product as an off-white solid (0.5 g, 35% yield). MS-ESI (*m*/*z*) $[M+H]^+$ 221.1.

To a solution of S20 (0.5 g, 2.27 mmol) in $CHCl_3$ (10 mL) was added bromine (116 µL, 2.27 mmol). The mixture was stirred at room temperature for 1 h. A colorless mixture was obtained. $CHCl_3$ was removed under vacuum. The crude product was purified by column chromatography (0 - 30% ethyl acetate in hexanes) to afford the product as a light yellow solid (0.4 g, 59% yield). MS-ESI (*m*/*z*) $[M+H]^+$ 299.0.

To a solution of S21 (0.4 g, 1.33 mmol) in acetonitrile (5 mL) was added 4-fluoro-2-aminopyridine (0.15 g, 1.33 mmol). The mixture was heated at 120 °C for 30 min. TLC and LC-MS showed completion of reaction. The acetonitrile was removed under vacuum and the crude product was purified by column chromatography (0 - 5% methanol in DCM) to afford the product as a light yellow solid (0.15 g, 36% yield). MS-ESI (*m*/*z*) $[M+H]^+$ 313.1.

To a solution of S22 (31 mg, 0.1 mmol) and piperazine (13 mg, 0.15 mmol) in DMSO was added $K_2CO_3$ (33 mg, 0.3 mmol). The reaction mixture was heated at 120 °C for 2 h. The mixture was cooled to room temperature, poured into ice water and filtered. The crude product was purified by silica gel column chromatography (0 – 10% $CH_3OH$ in $CH_2Cl_2$) to give **C30-Me** as a light yellow solid. (15 mg, yield 40%). HRMS-ESI (*m*/*z*) $[M+H]^+$ Calcd 379.1570; Found 379.1568.

$^1$H NMR (400 MHz, DMSO-$d_6$) δ 8.72 – 8.69 (m, 1H), 8.46 (s, 1H), 7.89 (d, $J$ = 9.0 Hz, 1H), 7.42 (dd, $J$ = 10.2, 2.5 Hz, 1H), 7.12 (dd, $J$ = 9.1, 2.4 Hz, 1H), 6.99 (td, $J$ = 7.6, 2.6 Hz, 1H), 6.90 (d, $J$ = 2.3 Hz, 1H), 3.40 (t, $J$ = 8.0 Hz, 4H), 2.93 (t, $J$ = 8.0 Hz, 4H), 2.76 (s, 3H).

$^{13}$C NMR (100 MHz, DMSO-$d_6$) δ 174.5, 165.0, 160.0 (d, $J$ = 247.7 Hz), 157.4, 155.0, 143.4 (d, $J$ = 14.2 Hz), 138.3, 129.1 (d, $J$ = 11.1 Hz), 126.8, 115.3, 114.8, 114.0, 113.4, 104.6 (d, $J$ = 29.4 Hz), 100.4 (d, $J$ = 23.5 Hz), 99.8, 47.3, 44.9, 20.6.

**3-(6-methoxyimidazo[1,2-*a*]pyridin-2-yl)-7-(piperazin-1-yl)-2*H*-chromen-2-one (C34b)**

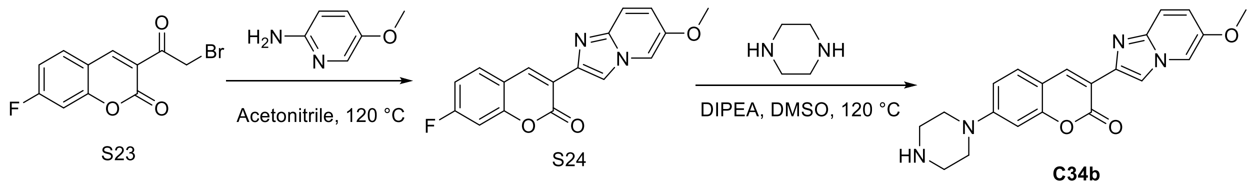

A mixture of 3-(2-bromoacetyl)-7-fluoro-2*H*-chromen-2-one (50 mg, 0.17 mmol) and 5-methoxy-2-aminopyridine (26 mg, 0.21 mmol) in acetonitrile was heated at 120 °C for 20 min in a Biotage Initiator+ microwave reactor. TLC and LC-MS showed completion of reaction. The reaction mixture was filtered, and the solid was washed with acetonitrile to afford a yellow solid which was used in the next step without further purification (41 mg, yield 74%). MS-ESI (*m*/*z*) $[M+H]^+$ 311.1.

To a solution of S24 (31 mg, 0.1 mmol) and piperazine (13 mg, 0.15 mmol) in DMSO was added $K_2CO_3$ (33 mg, 0.3 mmol). The reaction mixture was heated at 120 °C for 2 h. TLC and LC-MS showed completion of reaction. The mixture was cooled to room temperature, poured into ice water and filtered. The crude product was purified by silica gel column chromatography (0 – 10% $CH_3OH$ in $CH_2Cl_2$) to afford **C34b** as a yellow solid. (22 mg, yield 58%). HRMS-ESI (*m*/*z*) $[M+H]^+$ Calcd 377.1614; Found 377.1608.

$^1$H NMR (400 MHz, DMSO-$d_6$) δ 8.65 (s, 1H), 8.44 (s, 1H), 8.35 (d, *J* = 1.7 Hz, 1H), 7.67 (d, *J* = 8.8 Hz, 1H), 7.46 (d, *J* = 9.7 Hz, 1H), 7.06 (dd, *J* = 9.7, 2.4 Hz, 1H), 7.00 (dd, *J* = 8.9, 2.4 Hz, 1H), 6.85 (d, *J* = 2.4 Hz, 1H), 3.79 (s, 3H), 3.31 (t, *J* = 8.0 Hz, 4H), 2.85 (t, *J* = 8.0 Hz, 4H).

$^{13}$C NMR (100 MHz, DMSO-$d_6$) δ 160.1, 155.3, 154.1, 148.7, 141.9, 138.9, 138.4, 129.9, 120.9, 116.8, 115.4, 113.7, 112.1, 110.7, 109.5, 99.8, 56.6, 48.2, 45.6.

### NMR Spectra

SMSM6

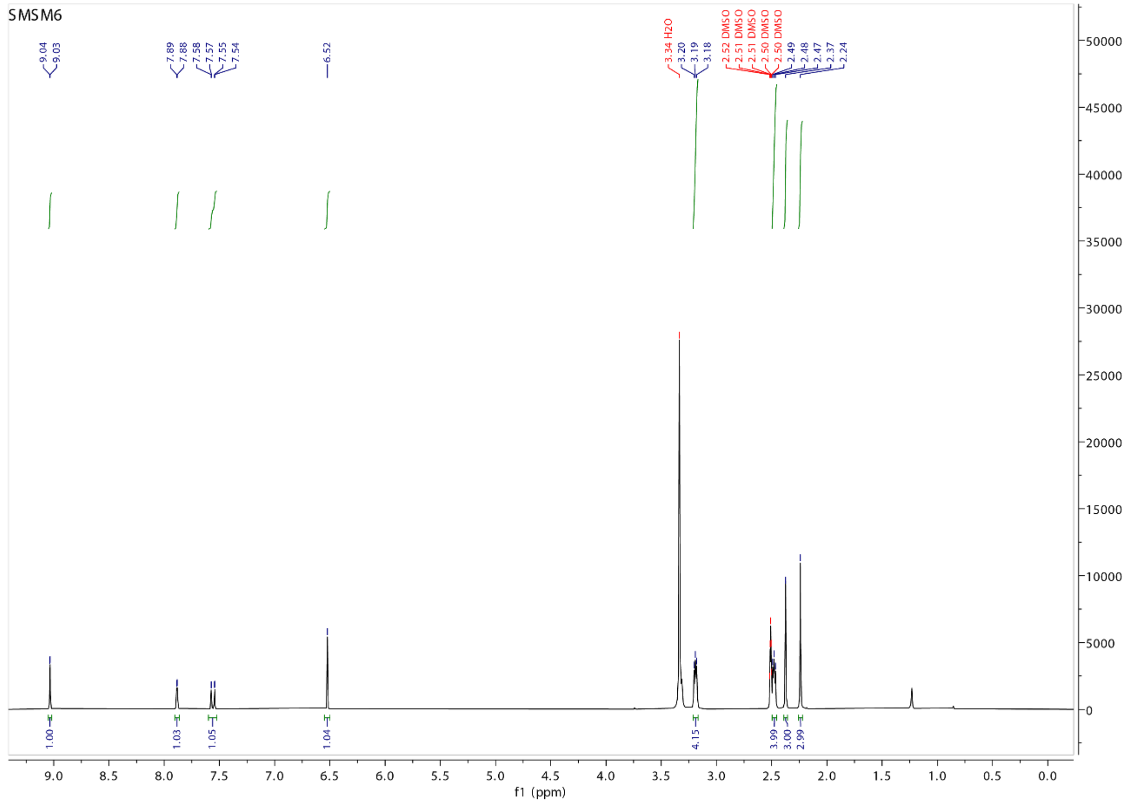

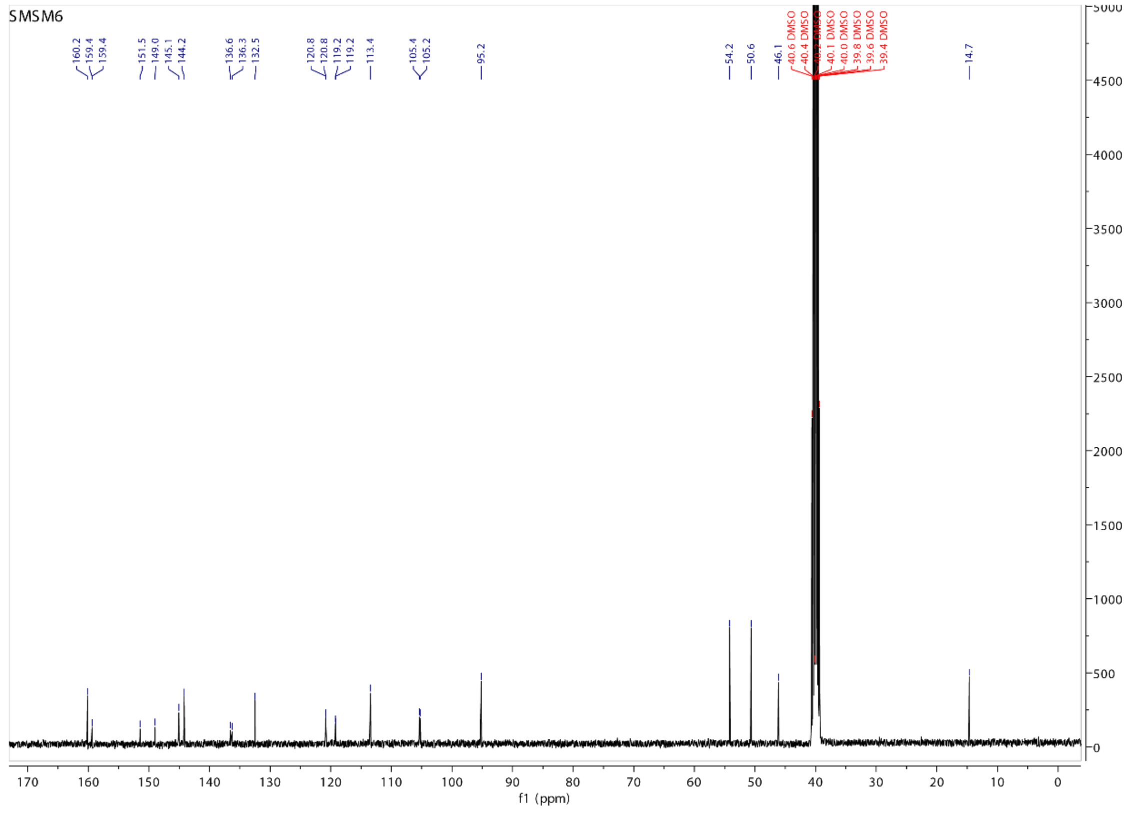

SMSM46

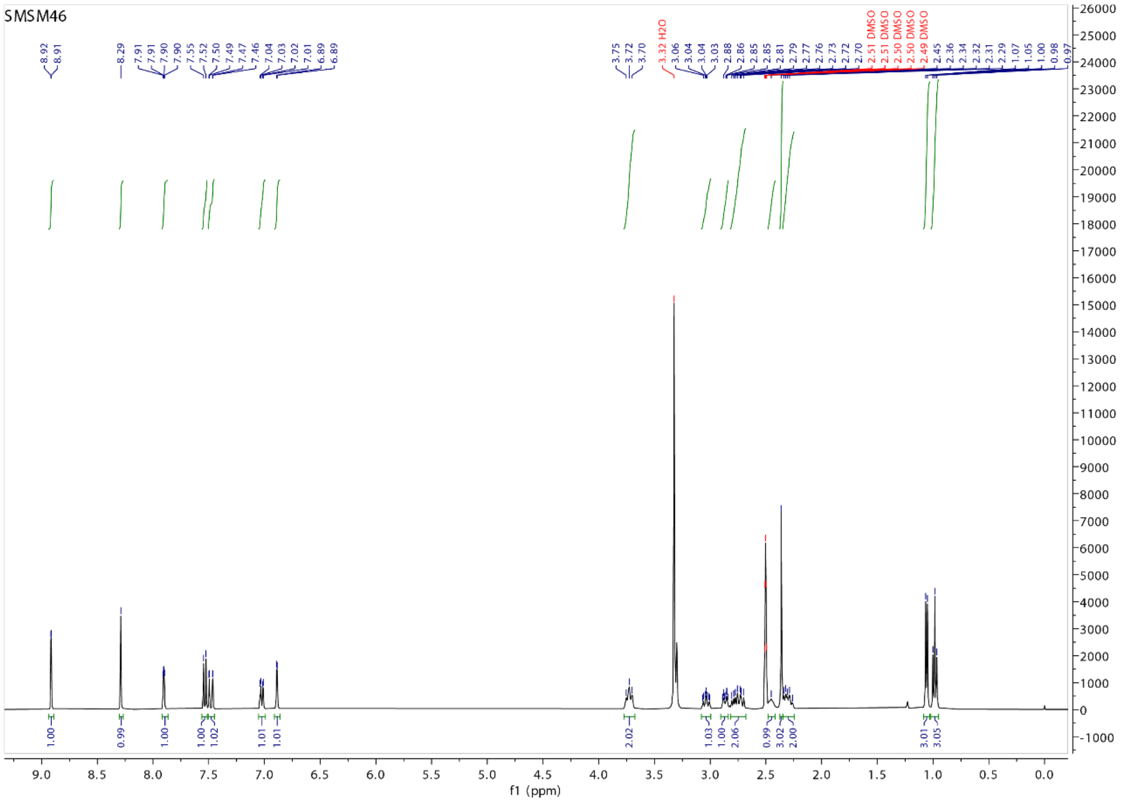

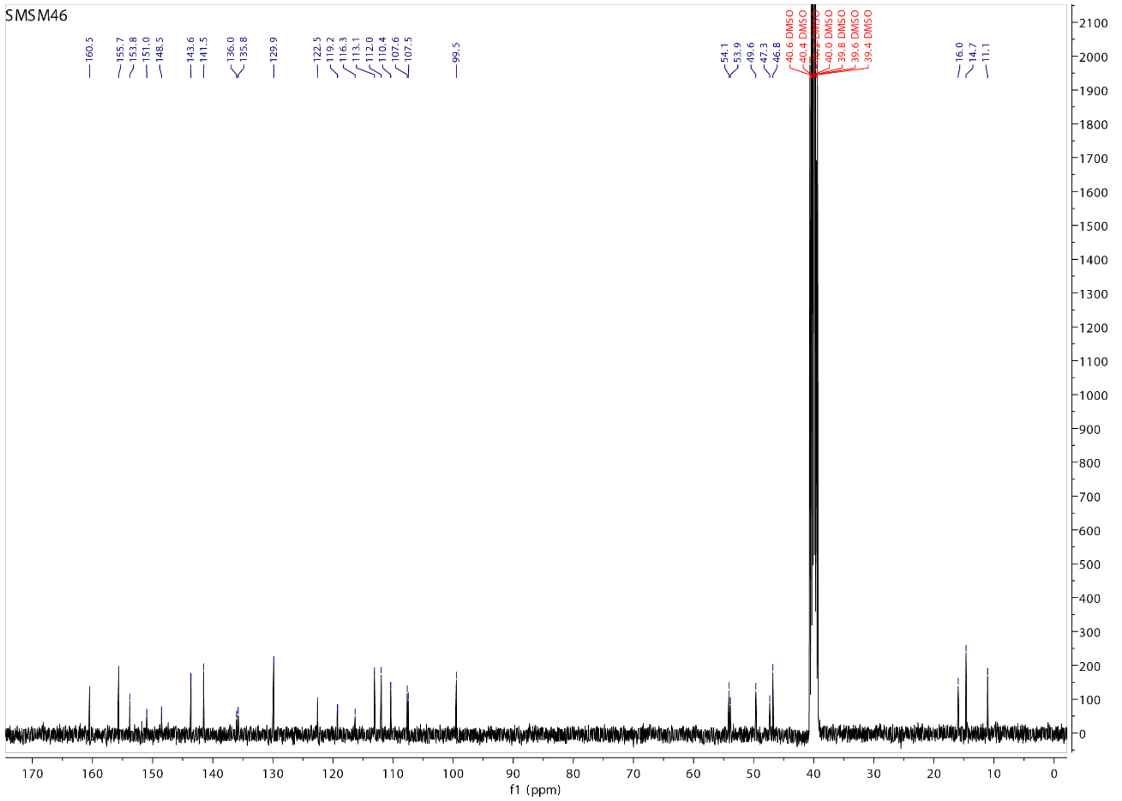

SMSM61

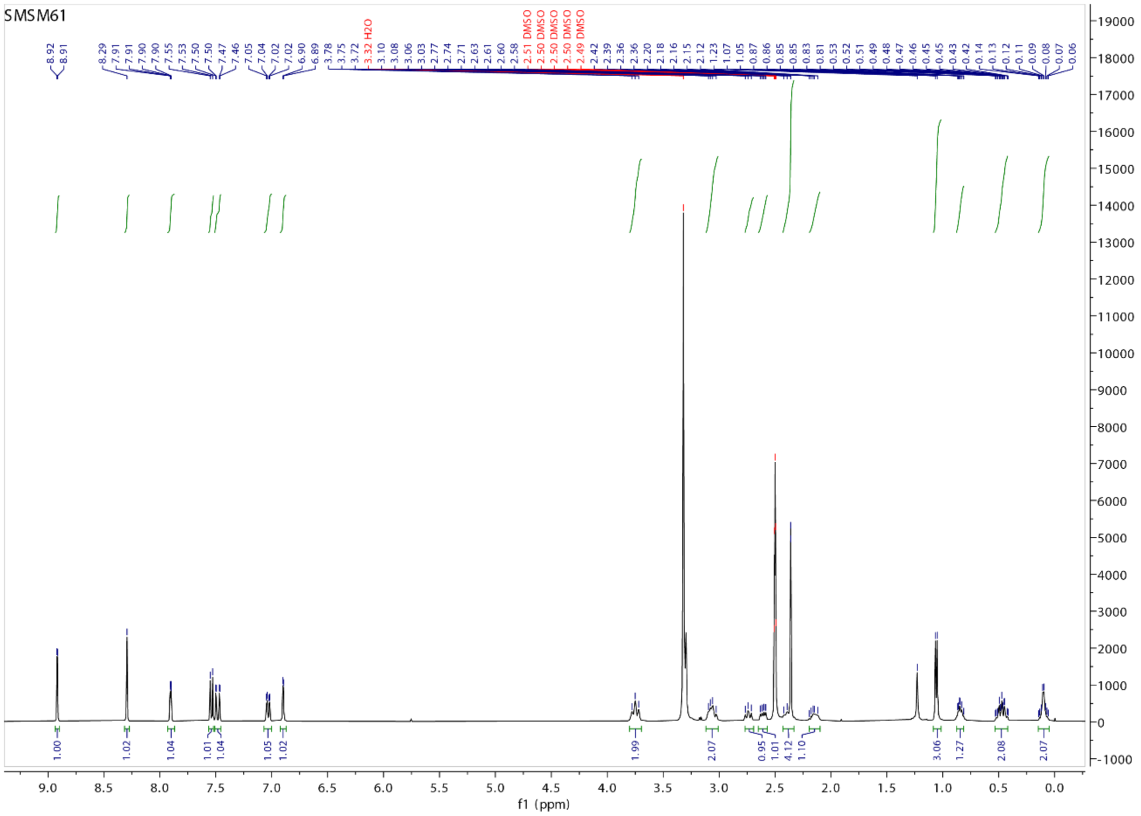
SMSM61
f1 (ppm)

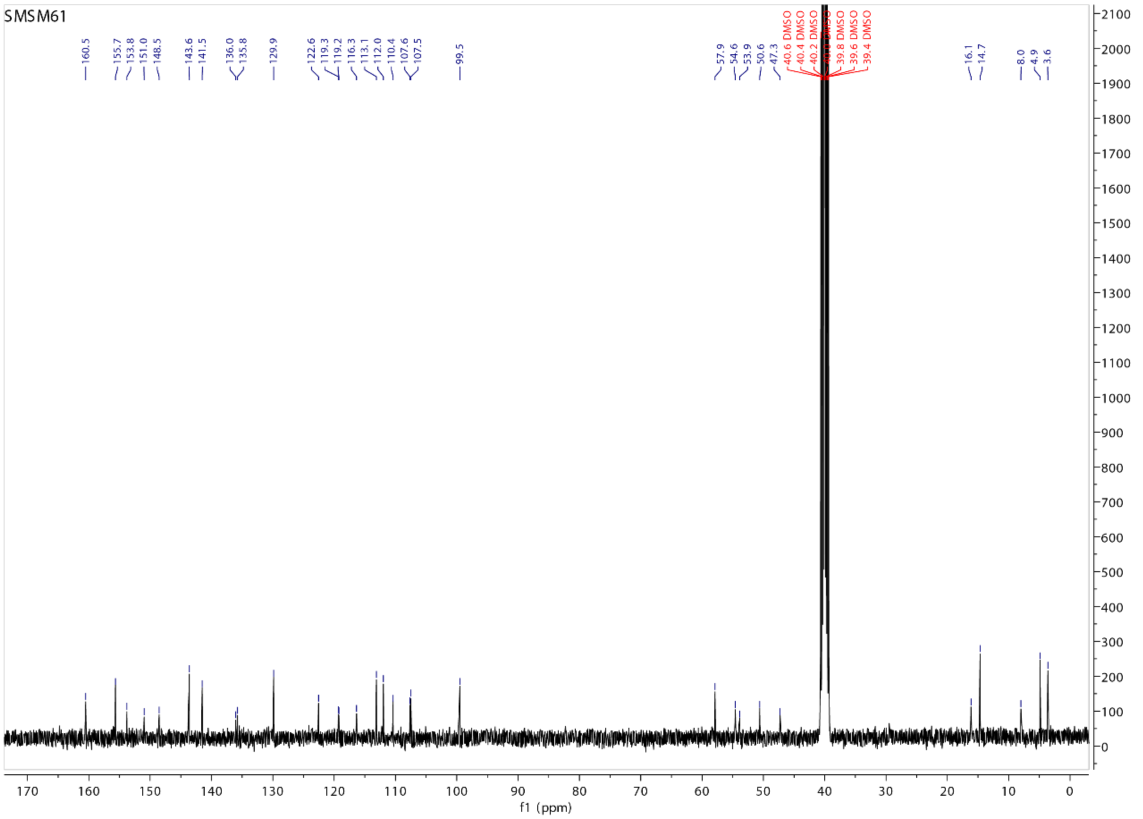
SMSM61
160.5
155.7
153.8
151.0
148.5
143.6
141.5
136.0
135.8
129.9
122.6
119.3
119.2
116.3
113.1
112.0
110.4
107.6
107.5
99.5
57.9
54.6
53.9
50.6
47.3
16.1
14.7
8.0
4.9
3.6
f1 (ppm)

SMSM64

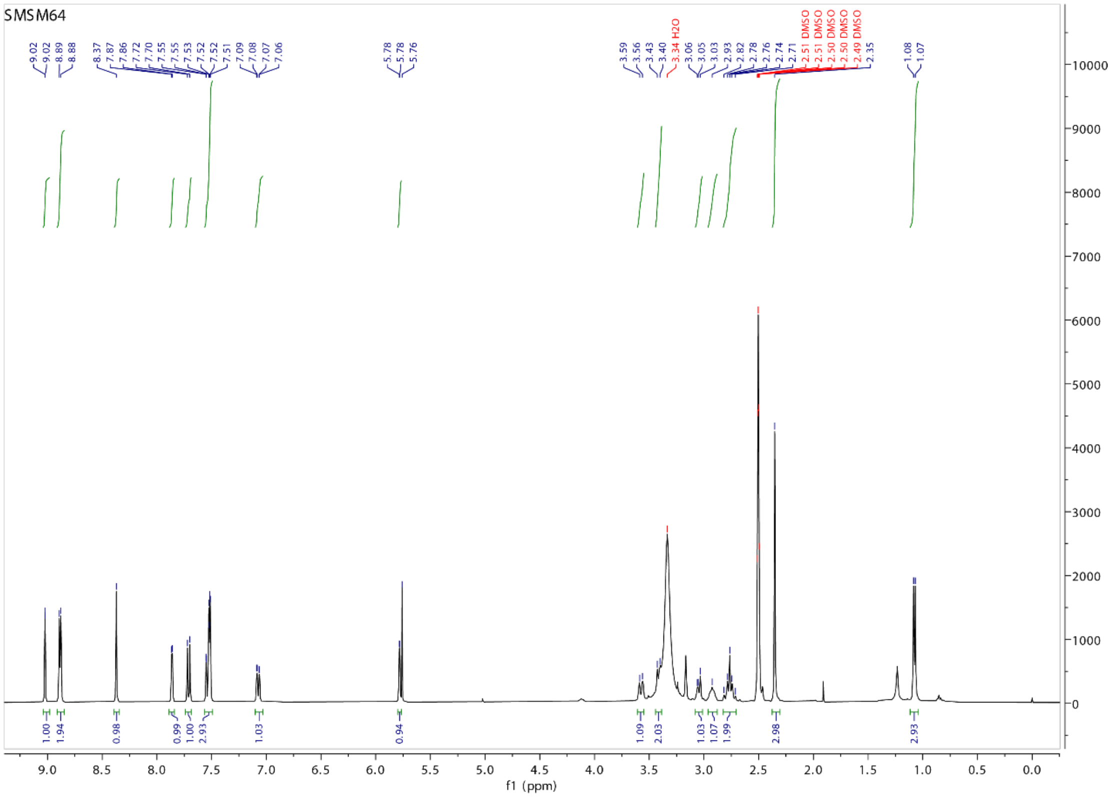
SMSM64
9.02
9.02
8.89
8.88
8.37
7.87
7.86
7.72
7.70
7.55
7.55
7.53
7.52
7.52
7.51
7.09
7.08
7.07
7.06
5.78
5.78
5.76
3.59
3.56
3.43
3.40
3.34 H2O
3.06
3.05
3.03
2.93
2.82
2.78
2.76
2.74
2.71
2.51 DMSO
2.51 DMSO
2.50 DMSO
2.50 DMSO
2.49 DMSO
2.35
1.08
1.07
1.00
1.94
0.98
0.99
1.00
2.93
1.03
0.94
1.09
2.03
1.03
1.07
1.99
2.98
2.93
f1 (ppm)

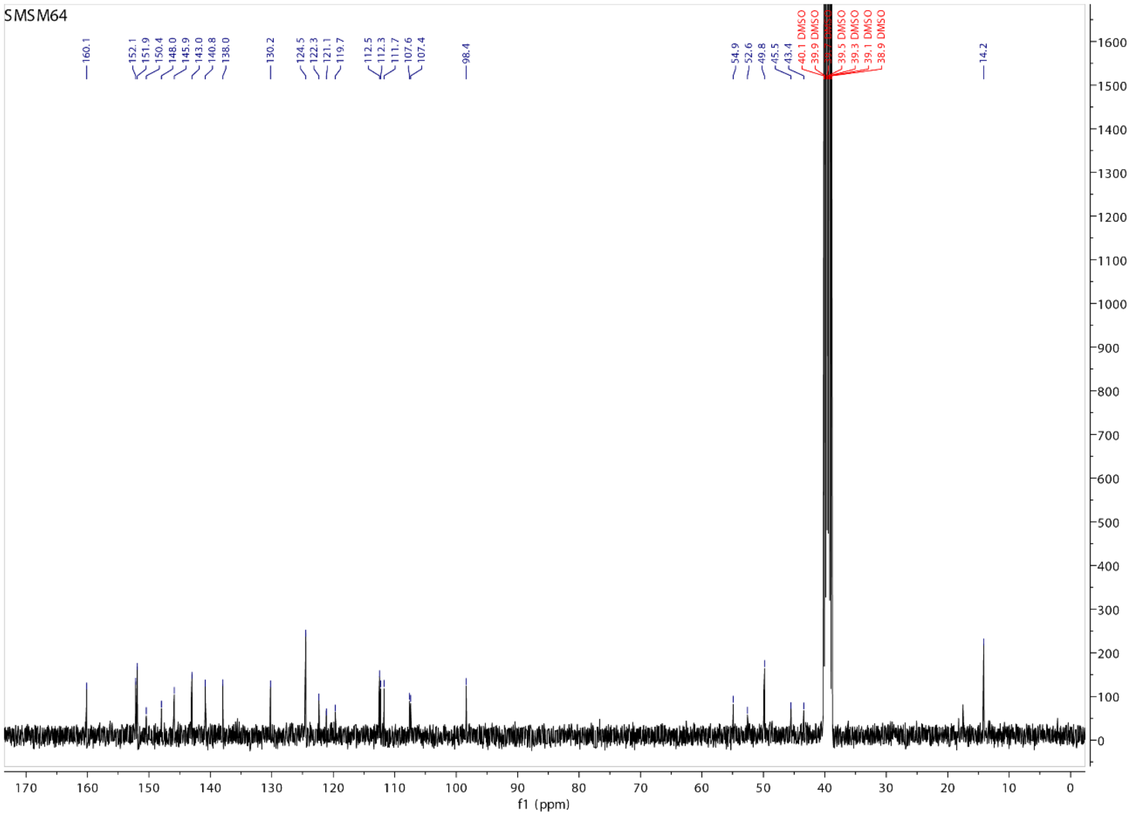
SMSM64
160.1
152.1
151.9
150.4
148.0
145.9
143.0
140.8
138.0
130.2
124.5
122.3
121.1
119.7
112.5
112.3
111.7
107.6
107.4
98.4
54.9
52.6
49.8
45.5
43.4
40.1 DMSO
39.9 DMSO
39.5 DMSO
39.3 DMSO
39.1 DMSO
38.9 DMSO
14.2
f1 (ppm)

C30-Ac

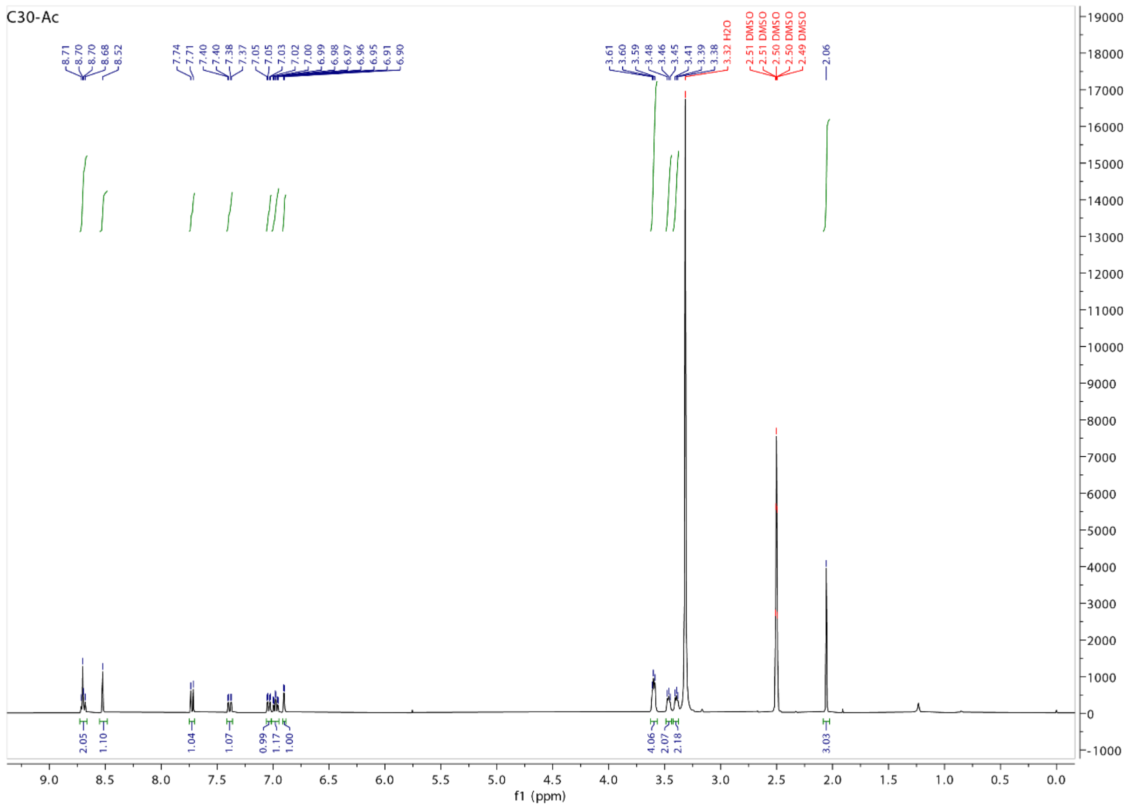

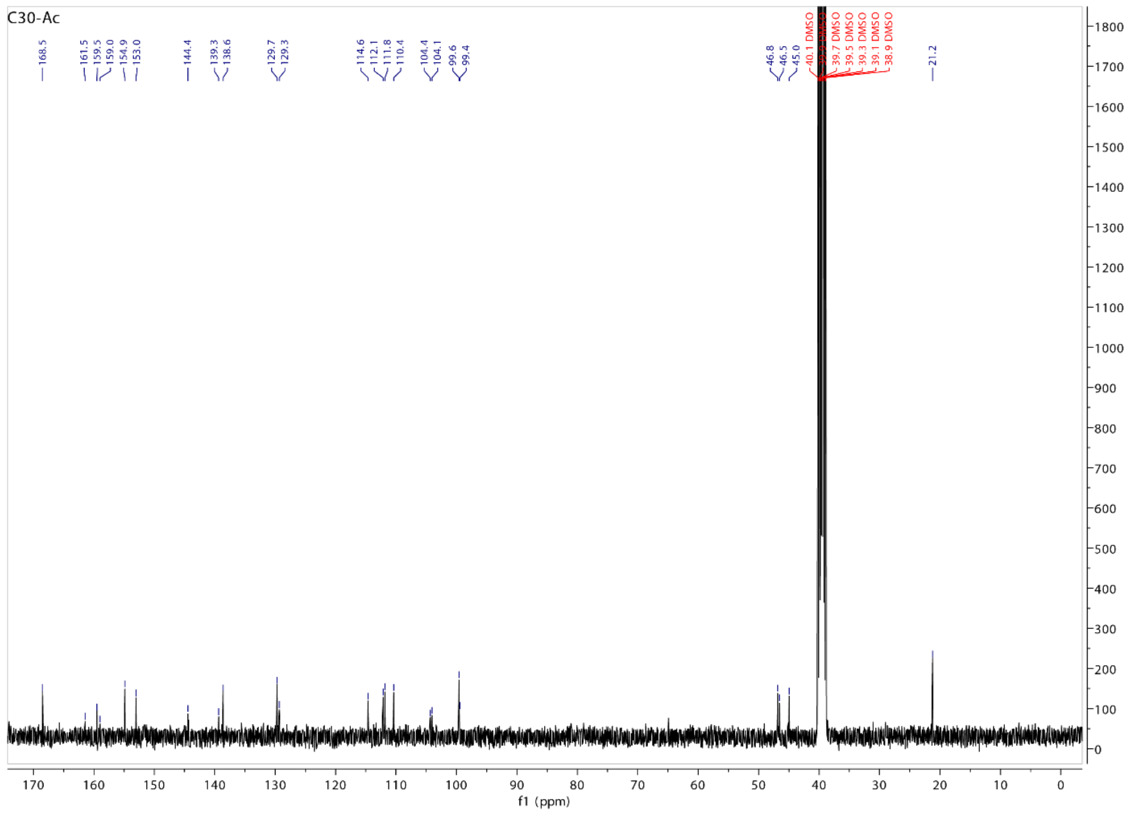

C30-Me$^{RingB}$

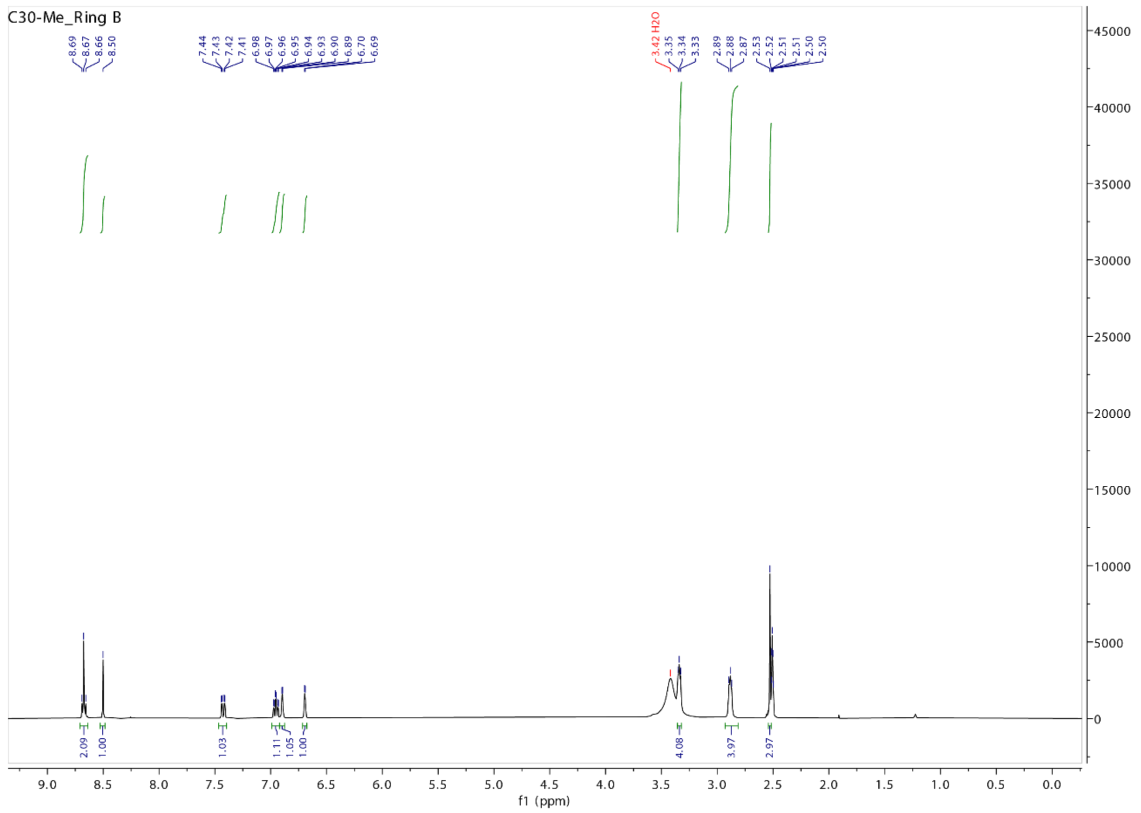
C30-Me_Ring B
8.69
8.67
8.66
8.50
7.44
7.43
7.42
7.41
6.98
6.97
6.96
6.95
6.94
6.93
6.90
6.89
6.70
6.69
3.42 H2O
3.35
3.34
3.33
2.89
2.88
2.87
2.53
2.52
2.51
2.51
2.50
2.50
2.09
1.00
1.03
1.11
1.05
1.00
4.08
3.97
2.97
f1 (ppm)

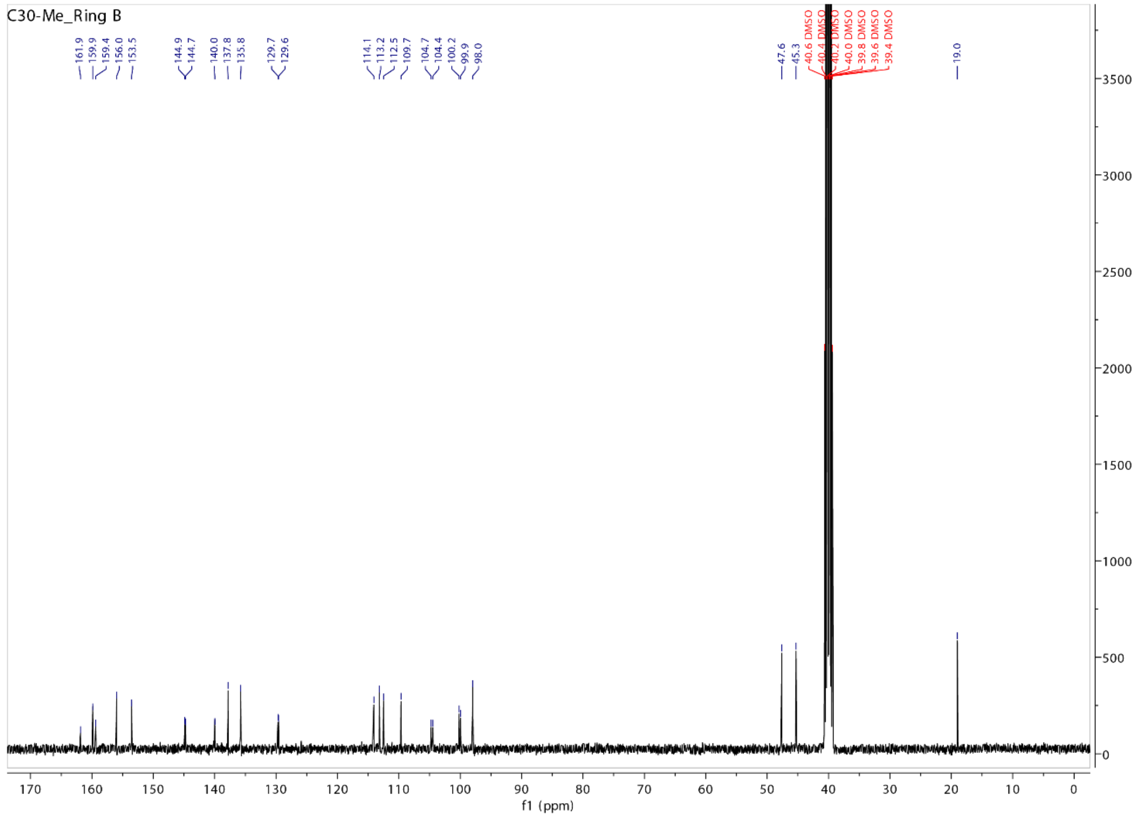
C30-Me_Ring B
161.9
159.9
159.4
156.0
153.5
144.9
144.7
140.0
137.8
135.8
129.7
129.6
114.1
113.2
112.5
109.7
104.7
104.4
100.2
99.9
98.0
47.6
45.3
40.6 DMSO
40.4 DMSO
40.2 DMSO
40.0 DMSO
39.8 DMSO
39.6 DMSO
39.4 DMSO
19.0
f1 (ppm)

C30-Me

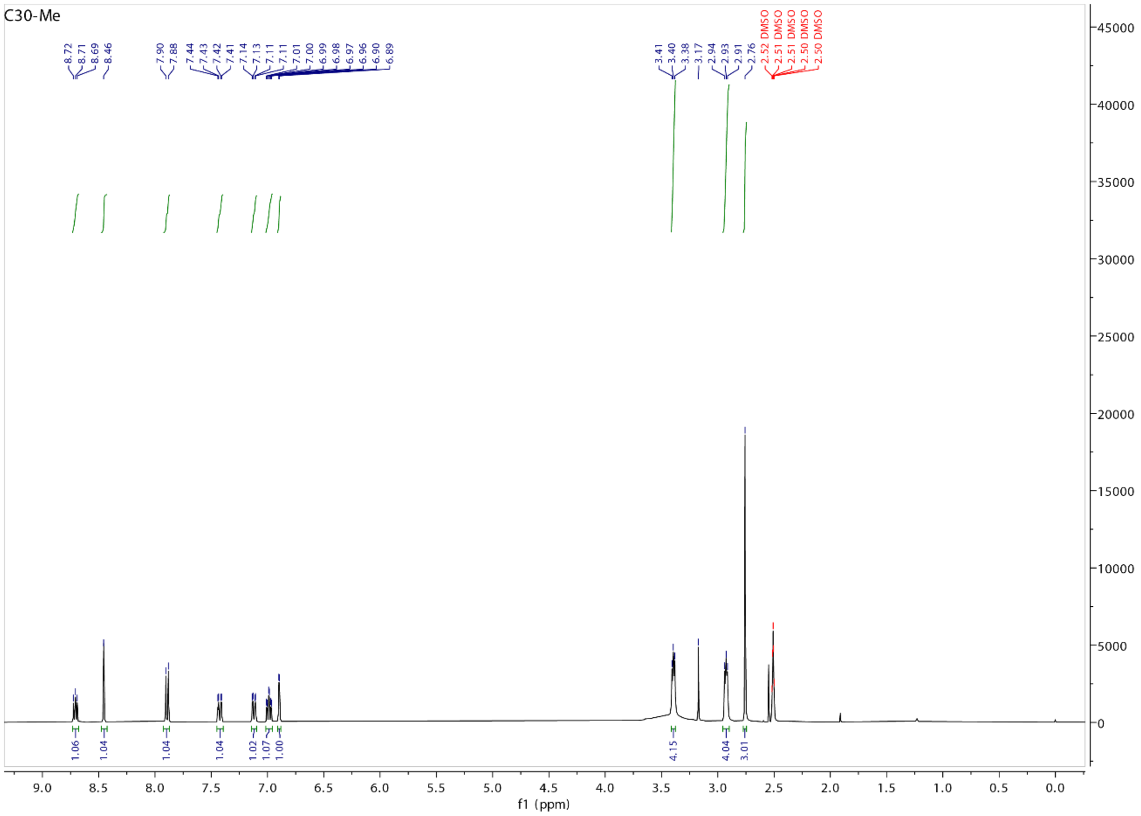

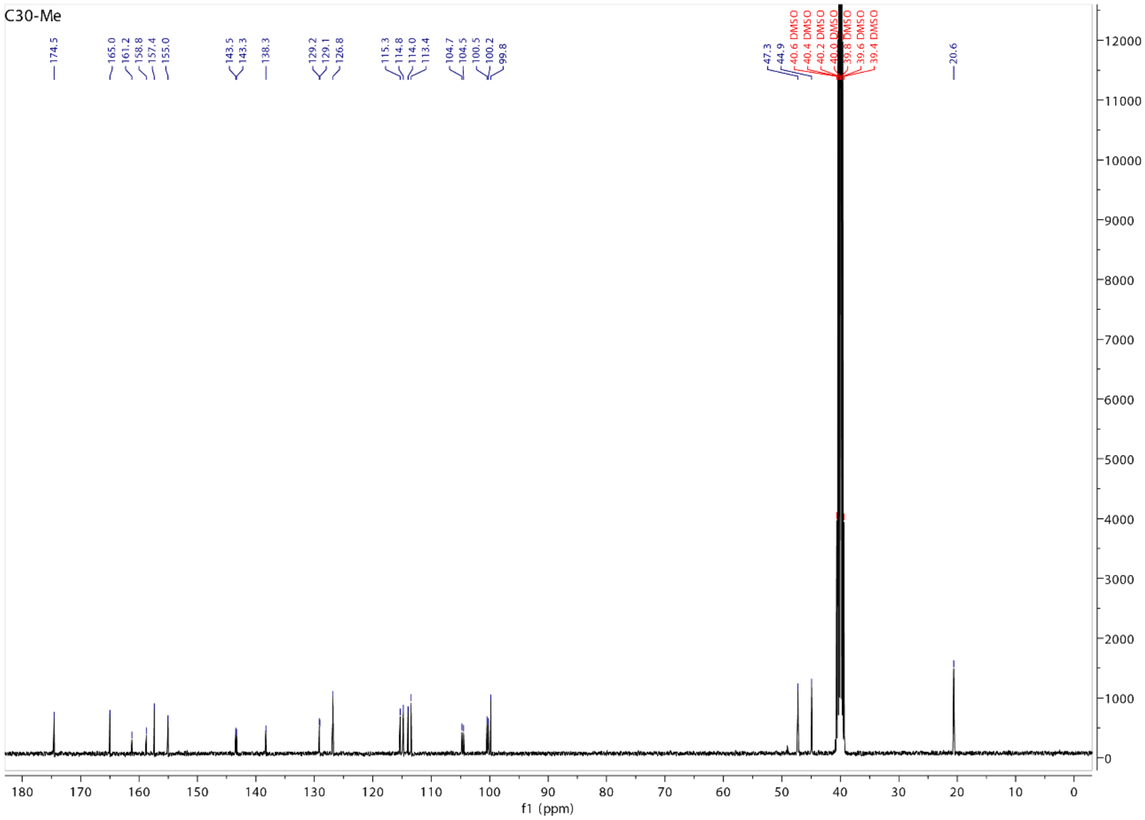

C34b

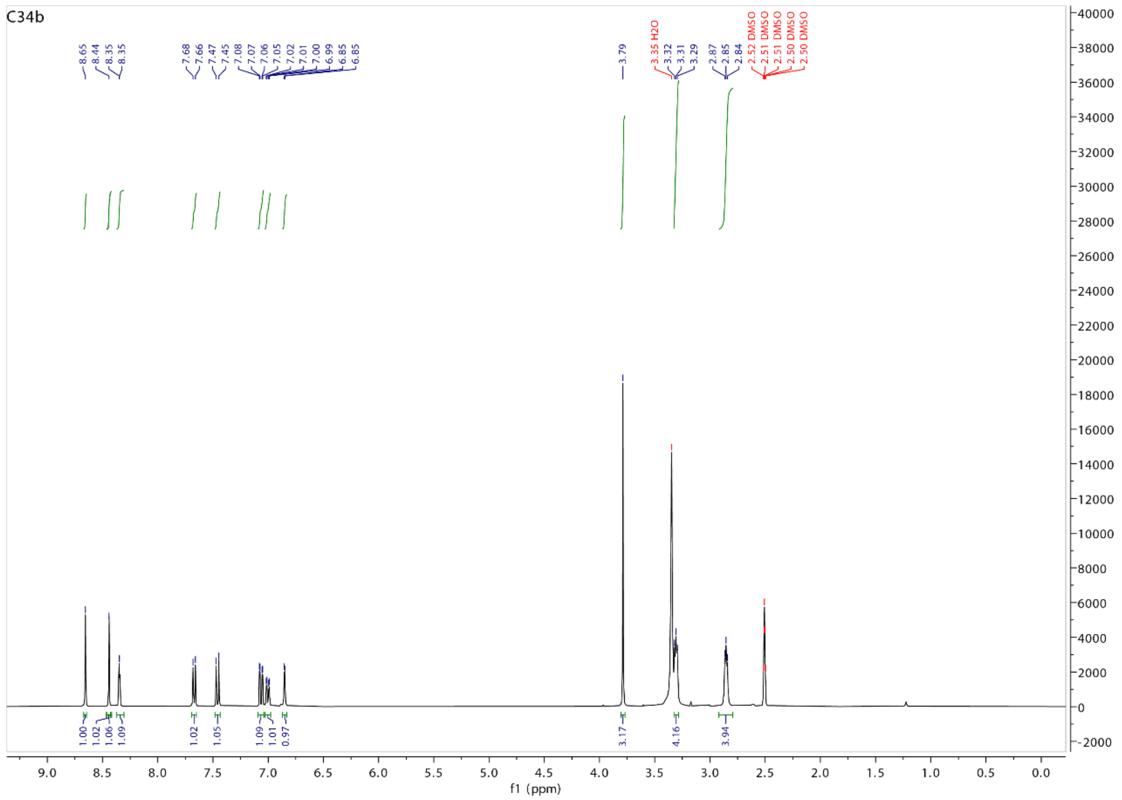
C34b
8.65
8.44
8.35
8.35
7.68
7.66
7.47
7.45
7.08
7.07
7.06
7.05
7.02
7.01
7.00
6.99
6.85
6.85
3.79
3.35 H2O
3.32
3.31
3.29
2.87
2.85
2.84
2.52 DMSO
2.51 DMSO
2.51 DMSO
2.50 DMSO
2.50 DMSO
1.00
1.02
1.06
1.09
1.02
1.05
1.09
1.01
0.97
3.17
4.16
3.94
f1 (ppm)

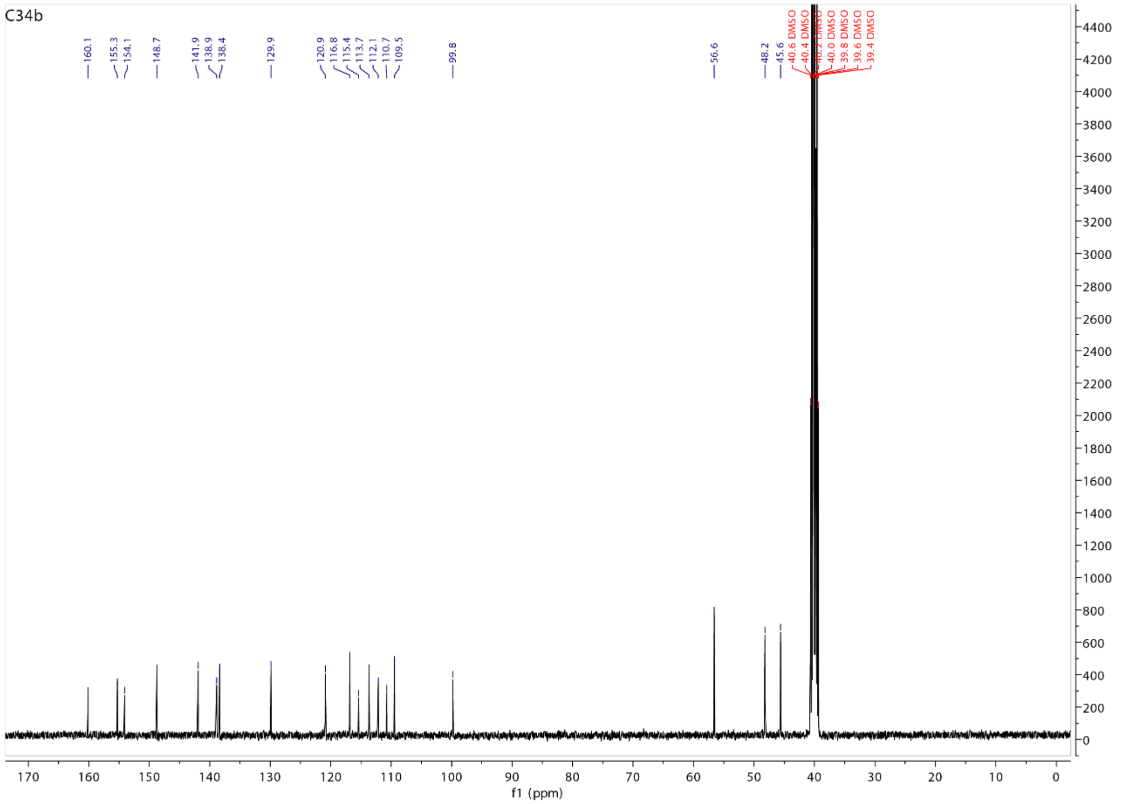
C34b
160.1
155.3
154.1
148.7
141.9
138.9
138.4
129.9
120.9
116.8
115.4
113.7
112.1
110.7
109.5
99.8
56.6
48.2
45.6
40.6 DMSO
40.4 DMSO
40.2 DMSO
40.0 DMSO
39.8 DMSO
39.6 DMSO
39.4 DMSO
f1 (ppm)
